# Obesogenic Diet Impairs Social Memory Through Alterations of Hippocampal CA2 Excitability and Oxytocin Signaling

**DOI:** 10.1101/2025.09.02.673707

**Authors:** Maud Muller, Alice Fermigier, Arielle Rakotonandrasana, Eva-Gunnel Ducourneau, Matteo Moschetta, Hazim Eldirdiri, Martina Radice, Pierre-Alexandre Raffaelli, Guillaume Ferreira, Vivien Chevaleyre, Rebecca A Piskorowski

## Abstract

While obesity induces cardio-metabolic disorders and cognitive deficits, the underlying neural mechanisms remain unexplored. In mice, exposure to an obesogenic high-fat and sugar diet (HFD) resulted in social recognition memory deficits in both males and females, a process that is dependent upon hippocampal area CA2 and oxytocin signaling. HFD-fed mice had stronger inputs onto CA2 pyramidal neurons that led to increased action potential firing, without altering intrinsic properties or inhibitory transmission. Chemogenetic CA2 pyramidal neuron inhibition in HFD-fed mice rescued social novelty discrimination, with novelty preference in males and familiarity preference in females. We also observed that HFD in both males and females impaired the ability of oxytocin receptor activation to be permissive for endocannabinoid-mediated plasticity. Furthermore, chemogenetic inhibition of CA2 pyramidal neurons rescued oxytocin-induced endocannabinoid plasticity in both HFD-fed male and female mice, whereas it prevented endocannabinoid plasticity in control diet-fed mice. In a concentration-dependent manner, oxytocin restored potentiation of excitatory responses, allowed for endocannabinoid plasticity at CA2 inhibitory synapses, and rescued social novelty discrimination in HFD-fed mice. This study uncovers mechanisms linking neuromodulation and endocannabinoid-mediated plasticity in social memory processes and reveals the deleterious consequences of an obesogenic diet on this process.

## INTRODUCTION

Nearly half of the global population and almost 20% of children and adolescents are overweight or obese. While these levels seem to be plateauing in high-income countries, global obesity levels are increasing in both young and adult populations^1,2^. Excessive caloric intake, particularly from western diets rich in saturated fat and refined sugar is a pivotal driver of this epidemic^3^. While the adverse metabolic consequences, cardiovascular risks, and the oncogenic potential of obesity are well-documented, the impact on cognitive function remains comparatively unexplored. Nonetheless, clinical studies have reported adverse effects of obesity or western diets on cognition and learning and memory performance^4–6^. With this evidence, the underlying mechanisms related to these changes are unexplored.

Amongst widespread impairments in cognition, rodents fed an obesogenic diet, have been reported to undergo hippocampal-dependent learning and memory deficits^7–10^. Interestingly, social recognition memory (SRM) has been shown to be impaired following exposure to an obesogenic high-fat and sugar diet (HFD)^11–15^. Area CA2 of the dorsal hippocampus (dCA2) has been shown by targeted lesions, chemogenetic and optogenetic studies to play a central role in SRM^16–19^. With this, afferents from the lateral entorhinal cortex (EC), hypothalamic paraventricular nucleus (PVN), supramammillary nucleus and septum have been proven necessary for SRM^20–23^. Interestingly, oxytocin (OT), a prosocial neuropeptide, seems to regulate dCA2 function. Specifically, dCA2 pyramidal neurons (PNs) express OT receptors (OTR), that, when activated result in increased excitability and spontaneous burst firing of CA2 PNs^24^ and also in a synaptic potentiation at the CA3-CA2 and EC-CA2 glutamatergic synapses^20,25^. OT inputs from the PVN have been shown to be important for SRM^20,26,27^. It is known though, that disruption in OT signaling is associated with obesity. OT plays a role in appetite suppression, has been shown to modulate food intake^28^ and studies have reported changes in plasma OT concentrations in obese individuals^28^. In HFD-induced obese mice, reduction of blood OT concentrations has been documented^29^, alongside a decrease in OT signaling in the hippocampus^14,30^. Additionally, the endocannabinoid (eCB) system has been highlighted as a key player in the pathogenesis of obesity^31^ and exposure to a HFD has been reported to alter endocannabinoid signaling in the hippocampus^32,33^. In area CA2, eCB-mediated inhibitory long-term depression (iLTD) of GABAergic inputs onto CA2 PNs has been shown to be required for SRM in standard diet-fed mice^34^. Thus, the exposure to HFD may be altering the function of CA2 by two separate mechanisms: eCB and OT signaling.

To understand the mechanisms underlying SRM deficits in HFD-fed mice, we examined synaptic transmission in area CA2, its OT signaling and eCB-mediated plasticity of GABAergic transmission. We provide evidence that the CA2 PNs of HFD-fed mice are hyperexcitable and chemogenetic inhibition of these cells results in a restoration of social novelty discrimination, with sex difference in novelty preference. Furthermore, we show that OT-induced potentiation of excitatory responses onto CA2 PNs is impaired in HFD-fed mice and that OT signaling in area CA2 is permissive for the eCB plasticity in control conditions but disrupted under HFD exposure. Additionally, chemogenetic inhibition can rescue OT-permissive eCB plasticity in HFD-fed mice. Higher concentrations of OT restored potentiation of excitatory responses and allowed for endocannabinoid dis-inhibitory eCB plasticity at CA2 synapses in slices prepared from HFD-fed mice. CA2 infusion of OT, in a concentration-dependent manner, rescued SRM deficits in HFD-fed mice.

## RESULTS

### HFD exposure induces SRM impairments

To determine how exposure to an obesogenic diet compromises physiology and hippocampal-dependent SRM, we exposed male and female mice to a HFD consisting of 40% kcal from saturated fat and 17% kcal from sugar for 12 weeks just after weaning at 21 days. This exposure lasted until mice were 15 weeks-old, an age that is considered to be adulthood^35^. This exposure to HFD induced significant weight gain and increased fat mass in both male and female mice (Sup Fig 1A-C). Shorter periods of exposure to the HFD of 10 days and 5 weeks did not result in significant weight gain.

To assess the effect of adolescent HFD exposure on SRM, we adapted a social discrimination test (from^36,37^ using 15 minutes and 3-hour delay between training and test (Fig 1A). To first examine sociability during training, we compared the exploration time of a novel social conspecific in a small cage, versus the same empty cage. Both groups explored more the social stimulus (Fig 1B) and exhibited a sociability index higher than 0 (Fig 1C; CD: n=31, one-sample t-test p=0.0001, HFD: n=25, one-sample t-test p=0.0001; CD vs HFD: unpaired t-test p=0.79) indicating normal sociability. Memory for the stimulus mouse was tested in both groups either 15 minutes or 3 hours later (Fig 1D-E). While both CD- and HFD-fed mice explored the novel stimulus mouse more than the familiar one after 15 minutes (CD: n=15,), only CD-fed mice showed similar performance after 3 hours (CD: n=16, one-sample t-test p=2.5E-3, HFD: n=15, one-sample t-test p=0.93; CD vs HFD: unpaired t-test p=0.024) whereas the mice exposed to a HFD did not discriminate between the familiar and novel mice. The fact that HFD-fed mice succeeded in discriminating between familiar and novel mouse after a 15 min delay supports the premise that there is a specific effect of HFD on memory.

**Figure 1.**
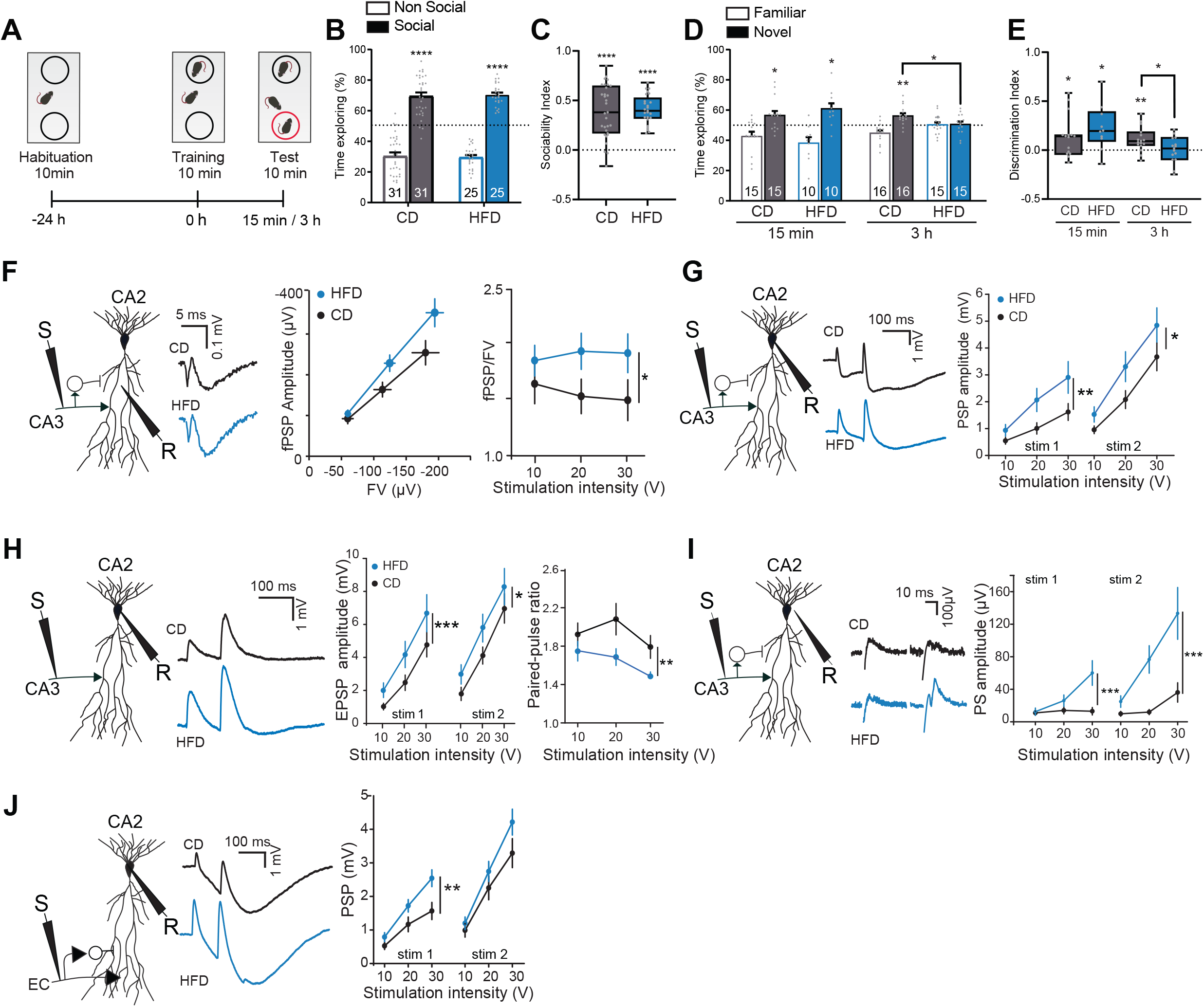
HFD exposure impairs social recognition memory and enhances area CA2 excitability. (A) Behavioral Paradigm. (B, C) Both CD-fed and HFD-fed mice preferred the social target as reflected by the percentage of time spent exploring the social stimulus compared to the non-social stimulus (CD: n=31, one-sample t-test p=0.0001, HFD: n=25, one-sample t-test p=0.0001; CD vs HFD: unpaired t-test p=0.79) as well as the sociability index. (D) After a 15-minute delay, both CD-fed and HFD-fed mice spent significantly more time exploring the novel mouse compared to the familiar one (CD: n=15, one-sample t-test p=2.4E-2, HFD: n=10, one-sample t-test p=0.013; CD vs HFD: unpaired t-test p=0.34). After a 3-hour delay, CD-fed mice still spent significantly more time exploring the novel mouse than the familiar one (CD: n=16, one-sample t-test p=2.5E-3, HFD: n=15, one-sample t-test p=0.93; CD vs HFD: unpaired t-test p=0.024), whereas HFD-fed mice spent equal amounts of time exploring both the novel and familiar mice leading to significant group difference. (E) The discrimination index was significantly greater than 0 in both CD-fed and HFD-fed mice after a 15-minute delay (CD: n=15, one-sample t-test p=2.4E-2, HFD: n=10, one-sample t-test p=0.013; CD vs HFD: unpaired t-test p=0.34). After a 3-hour delay, the discrimination index remained significantly greater than 0 for CD-fed mice (CD: n=16, one-sample t-test p=2.5E-3, HFD: n=15, one-sample t-test p=0.93; CD vs HFD: unpaired t-test p=0.024) but not for HFD-fed mice (HFD: n = 15, p > 0.05), with an index significantly higher in CD-fed mice compared to HFD-fed mice (p < 0.05). (F) Left: Schematic of recording configuration. Middle: Example traces of CA2 fPSPs upon SC stimulation for HFD and CD-fed mice. Right: fPSP vs FV graph for HFD and CD-fed mice at 10, 20 and 30V stimulation intensities; far right, HFD increases fPSP normalized on FV per stimulation intensity in CA2 upon SC stimulation (HFD: n=23,10, CD: n=13,6; Two-way ANOVA, p=0.016 for diet effect.). (G) Left: schematic of whole-cell recording configuration. Middle: example traces of PSPs recorded in HFD and CD-fed mice. Right: A HFD increases PSP amplitudes for a set stimulation intensity in CA2 PNs upon SC stimulation (HFD: n=34,18; CD: n=23,17; Two-way ANOVA, p=0.0052 for diet effect stimulation 1; p= 0.014 for diet effect stimulation 2). (H) Left: schematic of recording conditions. Middle: example EPSP traces in CA2 PNs upon SC stimulation. Right: A HFD increases EPSP amplitudes in CA2 PNs upon SC stimulation (CD: n=9,4; HFD: n=8,3; Two-way ANOVA, p=0.0064 for diet effect stimulation 1; p= 0.023 for diet effect stimulation 2); far right, a HFD decreases excitatory transmission paired-pulse ratio (CD: n=9,4; HFD: n=8,3; Two-way ANOVA, p=0.0041 for diet effect.). (I) Left: schematic of recording configuration. Middle: example population spike (PS) traces in CA2 upon SC stimulation. Right: A HFD enhances population spike amplitude upon SC stimulation (CD: n=18,7; HFD: n=23,6; Two-way ANOVA, p= 2.2E-4 for diet effect of stimulation 1; p=; 7.4E-6 for diet effect of stimulation 2). (J) Left: Schematic of recording conditions. Middle: example traces of transmission onto CA2 PNs upon PP stimulation. Right: A HFD increases PSP amplitude in CA2 PNs upon PP stimulation (CD: n=22,13; HFD: n=27,10; Two-way ANOVA, p= 0.0012 for diet effect of stimulation 1; p=0.099 for diet effect of stimulation 2).

### dCA2 excitability is enhanced following a HFD

As hippocampal area CA2 activity is required for SRM, we hypothesized that a HFD may induce alterations in this area that could lead to deficits in SRM. To test this, we used transverse hippocampal slices of HFD-fed and CD-fed mice and measured synaptic transmission in area CA2. Upon stimulation of Schaffer collateral (SC) afferents, field post-synaptic potentials recorded in stratum radiatum of area CA2 were larger in the HFD-fed group compared to the CD-fed group (Fig 1F, CD: n=13,6; HFD: n=23,10, Two-way ANOVA, p=0.2627), after normalizing to fiber volley (HFD: n=23,10, CD: n=13,6; Two-way ANOVA, p=0.016). We performed whole-cell recordings of CA2 PNs to directly examine changes in synaptic transmission at the CA3-CA2 synapse. In slices prepared from HFD-fed mice, we observed a consistent increase in the amplitude of post-synaptic potentials with SC stimulation (Fig 1G, HFD: n=34,18; CD: n=23,17; Two-way ANOVA, p=0.0052 for diet effect stimulation 1; p= 0.014 for diet effect stimulation 2).

To determine how much of this difference was due to inhibitory and excitatory transmission, we performed whole-cell voltage-clamp recordings to monitor inhibitory post synaptic potentials (IPSCs) at +10 mV both with and without AMPA and NMDA receptor blockers (NBQX and APV). No significant difference in the amplitude of IPSCs were observed between groups when we measured both feedforward inhibition (Sup Fig 2A, CD: n=18,11; HFD: n=17,12; Two-way ANOVA, p=0.086 for diet effect of stimulation 1; p=0.41 for diet effect of stimulation 2) or direct inhibition onto CA2 PNs (Sup Fig 2B, CD: n=18,11; HFD: n=17,12; Two-way ANOVA, p=0.40 for diet effect of stimulation 1; p= 0.12 for diet effect of stimulation 2). However, a decrease of the inhibitory paired-pulse ratio (PPR) of two subsequent IPSCs was observed in the HFD group (Sup Fig 2C, CD: n=18,11; HFD: n=17,12; Two-way ANOVA, p= 0.041 for diet effect), possibly indicating a change in presynaptic properties of GABA release in area CA2. Neither the amplitude not the frequency of spontaneous inhibitory activity onto area CA2 in the presence or absence of excitatory transmission blockers was altered by a HFD (Sup Fig 2D-G).

As inhibitory transmission onto CA2 was mostly unaltered by 12 weeks of a HFD, we measured the strength of excitatory transmission via patch-clamp recordings of CA2 PNs maintained at -70 mV in the presence of GABA_A_ and GABA_B_ receptor blockers (SR95531 and CGP55845). HFD exposure enhanced the strength of excitatory transmission onto CA2 PNs evoked by SC stimulation (Fig 1H, CD: n=9,4; HFD: n=8,3; Two-way ANOVA, p=0.0064 for diet effect stimulation 1; p=0.023 for diet effect stimulation 2). When we examined the ratio of the second evoked EPSP amplitude to the first, the ratio for the HFD-fed group was significantly lower as compared to the CD-fed group (Fig 1H, CD: n=9,4; HFD: n=8,3; Two-way ANOVA, p=0.0041 for diet effect). A reduction in PPR indicates that the probability of glutamate release at the CA3-CA2 synapse may be increased by 12 weeks of HFD.

To verify if the hyperexcitability of area CA2 influences the probability of CA2 PN action potential (AP) firing, we performed field recordings to record population spikes (PS) in the somatic layer of area CA2 upon SC stimulation. Stimulation of this input induced strong population-level AP firing of CA2 PNs in HFD-fed mice while stimulation of these inputs barely allowed for AP firing in control-diet mice, (Fig 1I, CD: n=18,7; HFD: n=23,6; Two-way ANOVA, p= 2.2E-4 for diet effect of stimulation 1; p= 7.4E-6 for diet effect of stimulation 2).

Because cortical input to area CA2 is also important for SRM^23^ and can strongly influence CA2 activity^38^, we next aimed to test whether the exposure to HFD also altered transmission at these synapses. Upon stimulation of EC afferents to CA2 in hippocampal slices, we observed that the amplitude of the PSP was significantly larger in HFD-fed mice (Fig 1J, CD: n=22,13; HFD: n=27,10; Two-way ANOVA, p= 0.0012 for diet effect of stimulation 1; p=0.099 for diet effect of stimulation 2). We saw no difference in inhibitory transmission with a stimulation pipette placed in stratum lacunosum moleculare (slm) (Sup Fig 3A, CD: n=8,6; HFD: n=5,4; Two-way ANOVA, p= 0.087 for diet effect of stimulation 1 without blockers of excitatory transmission; p= 0.093 for diet effect of stimulation 1 with blockers of excitatory transmission). However, we did detect a reduction in inhibitory current amplitude on the second pulse (Two-way ANOVA, p=1.2E-5 for diet effect of stimulation 2 without blockers of excitatory transmission; p=1.6E-5 for diet effect of stimulation 2 with blockers of excitatory transmission). Like what we observed at the CA3-CA2 synapse, we also detected a significant decrease in the PPR of inhibitory transmission when distal inhibitory inputs were directly stimulated (Sup Fig 3B, CD: n=8,6; HFD: n=5,4; Two-way ANOVA, p= 0.048 for diet effect), possibly indicating a change in the inhibitory transmission onto area CA2 in this dendritic compartment.

It has been reported that a single week of HFD exposure can have effects on hippocampal function^39^. Therefore, we examined synaptic transmission at different time points following exposure to HFD. We fed mice with a HFD or CD for 10 days or 5 weeks and compared the amplitude of the recorded CA3-CA2 field PSPs normalized by fiber volley amplitude. 10 days or 5 weeks of a HFD was not sufficient to increase SC to CA2 transmission (Sup Fig 4A-C, 10 days: CD: n=12,4; HFD: n=12,4, Two-way ANOVA, p=0.071; 5-weeks: CD: n=23,6; HFD: n=31,8, Two-way ANOVA, p=0.83). As HFD did not affect weight gain after 10 days or 5 weeks (Supp Fig 1B), this suggests HFD effect on CA2 excitability may require morphometric and metabolic changes as evidenced after 12 weeks of HFD exposure (Supp Fig 1B-C).

### HFD exposure does not alter CA2 PN intrinsic properties

To better understand the effect of HFD-exposure on CA2 PN physiology, we examined the intrinsic properties of the cells in both HFD-fed and CD-fed mice. HFD exposure did not affect CA2 PN resting membrane potential, membrane resistance, membrane capacitance, sag, rheobase current nor AP threshold (Sup Fig 5A-F). For a set injected current, CA2 PNs of HFD-fed mice fired APs similar to CD-fed CA2 PNs (Sup Fig 5G). Nevertheless, we observed that a HFD modified the AP shape of CA2 PNs upon current injection. Indeed, although rise time, amplitude and after-hyperpolarization of CA2 PN APs were unaltered, a HFD reduced the half-width and the decay-time constant of CA2 PN APs (Sup Fig 6A-F, CD: n=49,36; HFD: n=54,35; unpaired t-test, p=0.020 for AP half-width; p=0.035 for AP decay time-constant).

### Activation of hM4D(Gi) DREADDs in dCA2 PNs rescues social discrimination deficits in HFD-fed mice

We postulated that the compromised SRM we observed may be due to the increased synaptic transmission of dCA2 following exposure to a HFD, even though the intrinsic properties and synaptic plasticity were not altered. To test this hypothesis, we injected an AAV vector expressing floxed-hM4D(Gi) DREADDs or a control vector expressing floxed-mCherry bilaterally in the dCA2 area of Cacng5-Cre mice, expressing cre recombinase specifically in CA2^40^ to selectively inhibit CA2 PN activity following CNO administration (Fig 2A-B). After habituation to a test environment, CNO was administered in both CD and HFD-fed mice before a training trial when the mouse is presented a novel conspecific. We found that, in both male and female mice, sociability was unaltered by virus expression or diet (Fig 2C-D: CD-mCherry n=13, one-sample t-test p<0.0001; CD-Gi n=8, one-sample t-test p<0.0001; HFD-mCherry n=14, one-sample t-test p<0.0001; HFD-Gi n=11, one-sample t-test p<0.0001; Diet x Gi-DREADDs Two-way ANOVA F _(1, 42)_=0.024, p=0.88. G-H: CD-mCherry n=7, one-sample t-test p<0.0001; CD-Gi n=6, one-sample t-test p=5.00E-04; HFD-mCherry n=8, one-sample t-test p=2.60E-03; HFD-Gi n=7, one-sample t-test p=3.90E-03; Diet x Gi-DREADDs Two-way ANOVA (no effect) F _(1, 24)_=0.0007, p=0.98). The social discrimination test 3 hours later showed that CNO injection in the HFD-fed mice expressing hM4D(Gi) mice had a different effect in male and female mice. In male HFD-fed mice, hM4D(Gi) activation by CNO resulted in a preference for the novel over familiar conspecific, indicating optimal SRM (Fig 2E-F, CD-mCherry n=13, one-sample t-test p= 1.40E-03; CD-Gi n=8, one-sample t-test p=9.40E-01; HFD-mCherry n=14, one-sample t-test p=5.60E-01; HFD-Gi n=11, one-sample t-test p=0.0007; Diet x Gi-DREADDs Two-way ANOVA F _(1, 42)_=14.59, p=0.0004). In female HFD-fed mice, CA2 inhibition by hM4D(Gi) resulted in the mice showing a consistent preference for the familiar over the novel mouse (Fig 2 I-J, HFD Gi: n=7, p<0.01).CD-mCherry n=7, one-sample t-test p= 4.90E-03; CD-Gi n=6, one-sample t-test p=3.40E-01; HFD-mCherry n=8, one-sample t-test p=9.50E-01; HFD-Gi n=7, one-sample t-test p=0.0013; Diet x Gi-DREADDs Two-way ANOVA (no effect) F _(1, 24)_=16.19, p=0.0005). Thus, the ability to discriminate between novel and familiar conspecifics is restored by CA2 inhibition in both sexes despite being based on opposite novelty/familiarity preference in HFD-fed males and females.

**Figure 2.**
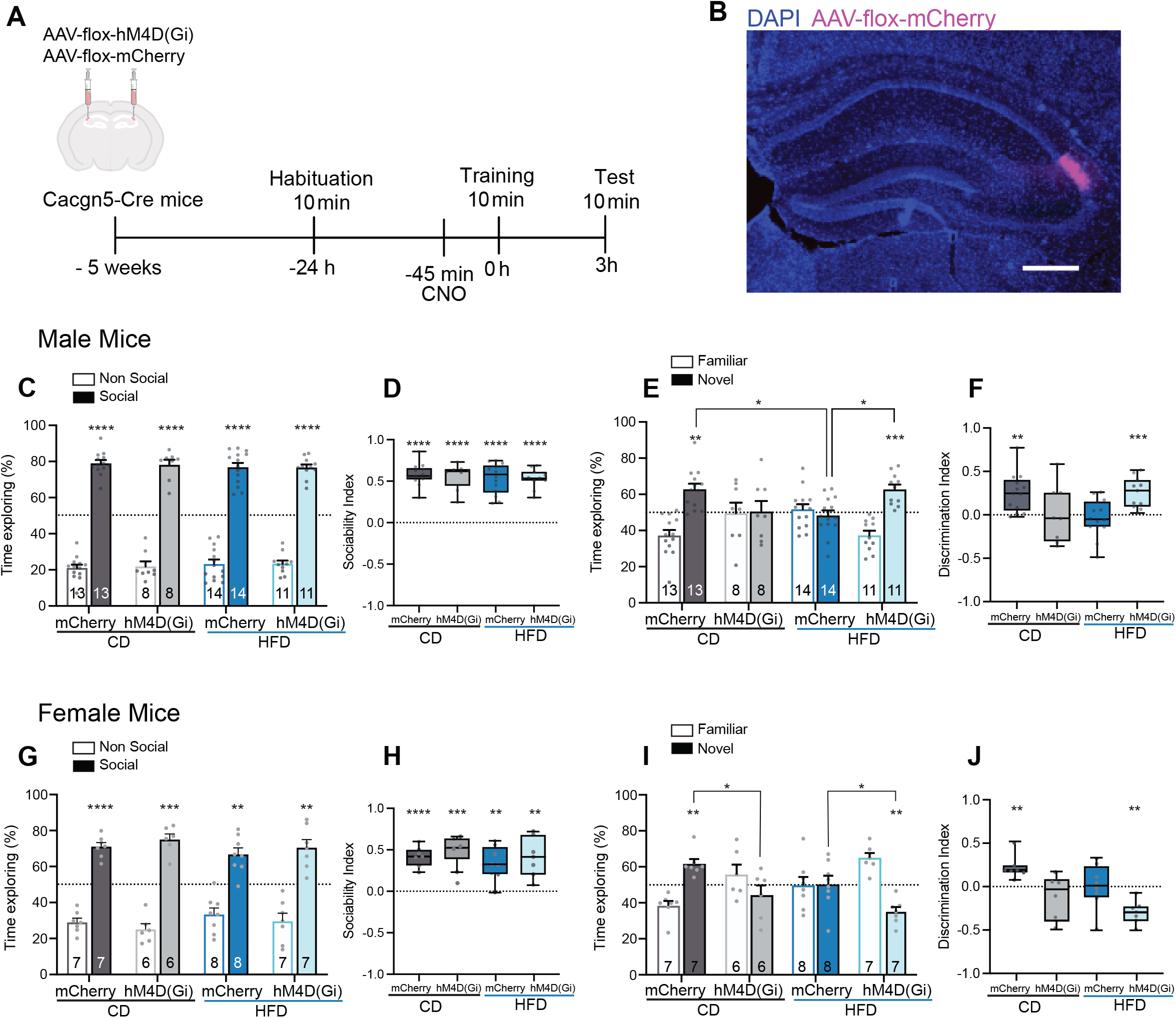
HFD-induced social memory deficits can be rescued by activation of Gi-DREADD in CA2 PNs. (A) Behavioral paradigm of AAV-mediated Gi-DREADD expression in CA2 PNs for SRM testing. (B) Example image of AAV virus-mediated delivery of vectors in dCA2. Scale bar is 200 µm. (C) HFD and CD male mice, injected with either mCherry control virus or Gi-DREADD virus, explore the social stimulus a greater amount of time compared to the non-social one to 50%). (D) Sociability index for social versus non-social stimuli is different than 0 in both CD and HFD-fed male mice (CD-mCherry n=13, one-sample t-test p<0.0001; CD-Gi n=8, one-sample t-test p<0.0001; HFD-mCherry n=14, one-sample t-test p<0.0001; HFD-Gi n=11, one-sample t-test p<0.0001; Diet x Gi-DREADDs Two-way ANOVA F _(1, 42)_=0.024, p=0.88). (E) In the test phase, CD-mCh male mice explore the novel mouse a greater amount of time compared to the familiar one. CD-Gi and HFD-mCh male mice explore the novel and familiar mice the same amount of time. HFD-Gi male mice prefer exploring the novel mouse over the familiar one. Percentage time exploring the novel mouse is greater in CD-mCh males and HFD-Gi vs HFD-mCh males (Two-way ANOVA, interaction Diet x Gi-DREADDs followed by post-hoc tests, p=0.0004 for both comparisons). (F) Discrimination index is higher than 0 in CD-mCh and HFD-Gi, but not in CD-Gi or HFD-mCh male mice (CD-mCherry n=13, one-sample t-test p= 1.40E-3; CD-Gi n=8, one-sample t-test p=9.40E-1; HFD-mCherry n=14, one-sample t-test p=5.60E-1; HFD-Gi n=11, one-sample t-test p=0.0007; Diet x Gi-DREADDs Two-way ANOVA F _(1,42)_=14.59, p=0.0004). (G) HFD and CD female mice, injected with either mCherry control virus or Gi-DREADD virus, explore more the social stimulus than the non-social one. (H) Sociability index for social versus non-social stimuli is different than 0 in both CD and HFD-fed female mice (CD-mCherry n=7, one-sample t-test p<0.0001; CD-Gi n=6, one-sample t-test p=5.00E-4; HFD-mCherry n=8, one-sample t-test p=2.60E-3; HFD-Gi n=7, one-sample t-test p=3.90E-3; Diet x Gi-DREADDs Two-way ANOVA (no effect) F _(1, 24)_=0.0007, p=0.98). (I) In the test phase, CD-mCh female mice explore the novel mouse a greater amount of time compared to the familiar one. CD-Gi and HFD-mCh female mice explore the novel and familiar mice the same amount of time. HFD-Gi female mice explore the familiar mouse a greater amount of time compared to the novel one. Percentage time exploring the novel mouse is greater in CD-mCh males than CD-Gi and less in HFD-Gi than HFD-mCh females. (J) Discrimination index is higher than 0 in CD-mCh females and less in HFD-Gi females, but not in CD-Gi or HFD-mCh male mice (CD-mCherry n=7, one-sample t-test p= 4.90E-3; CD-Gi n=6, one-sample t-test p=3.40E-1; HFD-mCherry n=8, one-sample t-test p=9.50E-1; HFD-Gi n=7, one-sample t-test p=0.0013; Diet x Gi-DREADDs Two-way ANOVA (Gi-DREADD effect) F _(1, 24)_=16.19, p=0.0005). The number of animals used per group is shown on the figure.

Interestingly, HFD or CA2 inactivation did not affect social recognition when the mice were discriminating between a novel mouse or a familiar cage mate, as all groups investigated the novel mouse, indicating an effect of HFD and CA2 manipulation on novel social memory formation rather than on social discrimination *per se* (Sup Fig 7A-C). Moreover, inhibition of CA2 PNs did not rescue HFD-induced deficits in object recognition memory (Sup Fig 7D-F), a task that does not require area CA2^16^.

### HFD exposure results in altered oxytocin receptor signaling in area CA2

Following exposure to a HFD, a reduction in hippocampal OT signaling has previously been reported^14,30^. To examine this further in area CA2, we measured OTR expression in HFD-fed or CD-fed mice in hippocampal areas CA1, CA2 and the dentate gyrus at baseline (home cage) or following a novel social interaction. In area CA1, there was a significant effect of diet on OTR gene expression with lower expression in HFD-fed mice for both the novel social and control conditions (Fig 3A). Interestingly, in area CA2 there was a significant interaction between diet and condition. We detected a significant increase in OTR gene expression in CD-fed mice that had encountered a novel conspecific as compared to home-cage controls and to HFD-fed mice that had social interaction. In HFD-fed mice, however, there was no significant difference in the OTR gene expression in mice exposed to a novel conspecific as compared to home-cage controls (Fig 3A). For the dentate gyrus, we detected no effect of diet, condition or interaction between diet and condition on OTR gene expression.

**Figure 3.**
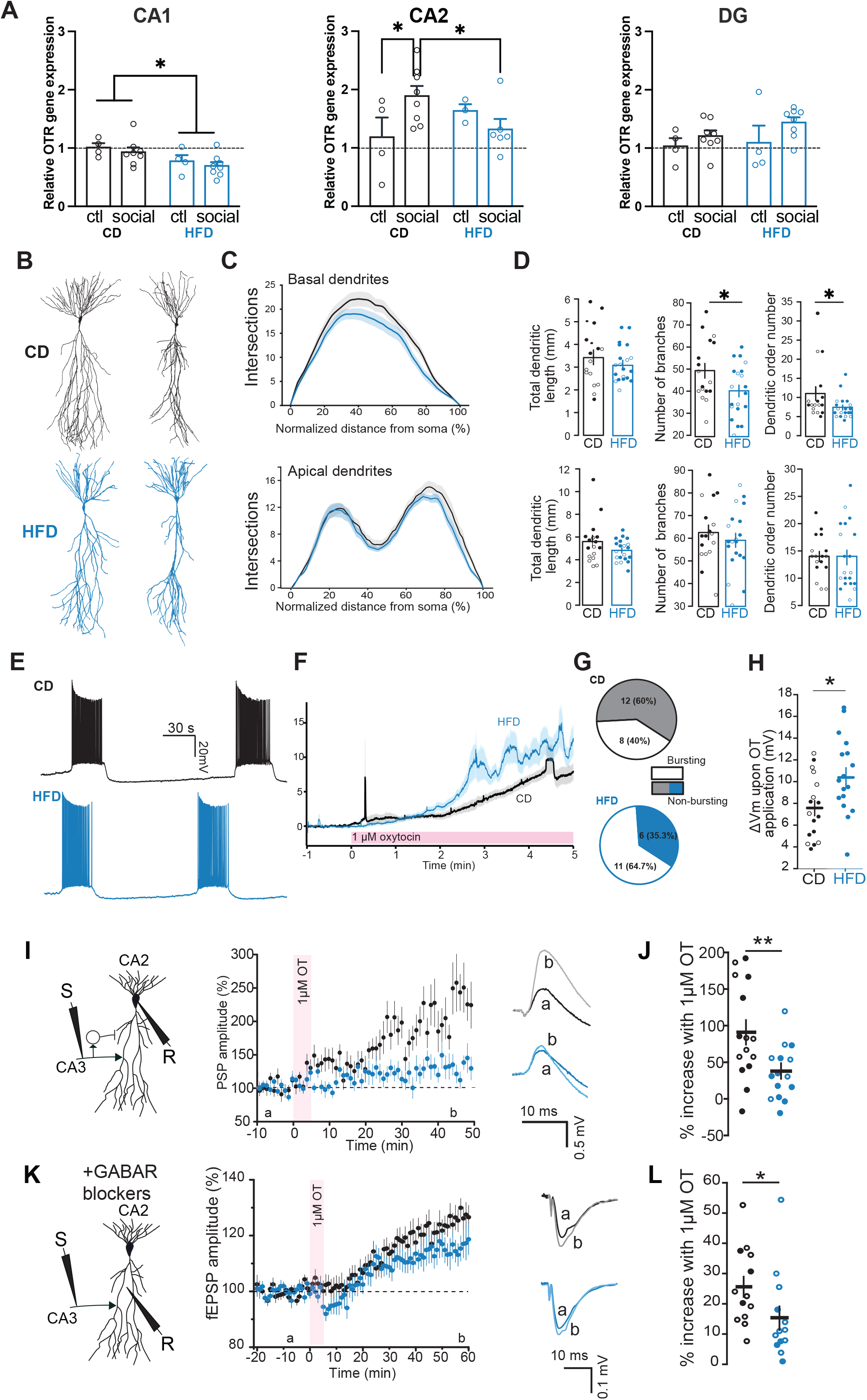
A HFD reduces OTR gene expression, dendritic complexity and OT-induced long-term potentiation of CA3-CA2 responses. (A) OTR gene expression, as detected by qPCR, in areas CA1, CA2 and dentate gyrus in mice 60 minutes following exposure to either a novel mouse for 10 minutes (social, n=8) or their home cage (ctl, n=4) for both CD and HFD conditions (CA1: Two-way ANOVA, diet effect p<0.01; CA2: Two-way ANOVA p<0.01; DG: Two-way ANOVA p>0.05). (B) Example reconstructions of CA2 PNs of CD (top) and HFD (bottom, blue) fed mice. (C) Sholl analysis of basal (top) and apical (bottom) dendrites for HFD (blue) and CD (black) CA2 PNs (basal: CD: n=17,10; HFD: n=20,13) (apical: CD: n=17,10; HFD: n=20,13). (D) Top left, total basal dendritic length for CD and HFD CA2 PNs (CD: n=17,10; HFD: n=20,13; unpaired t-test p=0.46). Top center, number of basal branches for CD and HFD CA2 PNs (CD: n=17,10; HFD: n=20,13; unpaired t-test p=0.036) Top right, number of orders of basal dendrites for CD and HFD CA2 PNs (CD: n=17,10; HFD: n=20,13; unpaired t-test p=0.044). Bottom left, total apical dendritic length for CD and HFD CA2 PNs (CD: n=17,10; HFD: n=20,13; unpaired t-test p=0.12). Bottom center, number of apical branches for CD and HFD CA2 PNs (CD: n=17,10; HFD: n=20,13; unpaired t-test p=0.39). Bottom right, number of orders of apical dendrites for CD and HFD CA2 PNs (CD: n=17,10; HFD: n=20,13; Mann-Whitney U test p=0.60). Filled circles represent CA2 PNs from male mice; empty circles represent CA2 PNs from female mice. (E) Example traces of OT-induced CA2 PN bursting for CA2 PNs from CD and HFD fed mice. (F) Average depolarization of CA2 PN upon 1µM OT application from 17 neurons each from CD and HFD-fed mice. (G) Top, 8 of 20 (40%) of CA2 PNs of CD-fed mice burst upon 1µM OT application. Bottom: 11 of 17 (64.7%) of CA2 PNs of HFD-fed mice burst upon 1µM OT application (CD: n=20,9; HFD: n=17,9; χ-square test, p=0.13). (H) OT-induced depolarization 5-minutes after application onset is stronger in CA2 PNs of HFD compared to CD-fed mice (CD: n=17,9, 7.56±0.72mV; HFD: n=17,9, 10.56±0.86mV; Mann-Whitney U test p=0.018). (I) Left: Schematic of whole-cell PSP recording of CA2 PN upon SC stimulation. Middle: 5-minutes 1µM OT application induces a stronger long-term increase in PSP amplitude in CA2 PNs in CD versus HFD-fed mice. Right: Example traces of 1µM OT-induced CA2 PSP responses in CD and HFD-fed mice. (J) Percentage of 1µM OT-induced PSP potentiation for each recorded CA2 PN (measured at 10 last minutes of recording normalized to baseline) (CD: n=15,13 one-sample t-test p=1.05E-4 baseline vs end; HFD: n=15,12 one-sample t-test p=0.0023 baseline vs end; CD vs HFD: unpaired t-test p=0.0081). Filled circles represent CA2 PNs from male mice; empty circles represent CA2 PNs from female mice. (K) Left: Schematic of field EPSP recording of CA2 PN upon SC stimulation in the presence of blockers of inhibitory transmission. Middle: 5-minutes 1µM OT application induces a stronger long-term increase in CA3-CA2 fEPSP amplitude in CD compared to HFD-fed mice. Right: Example traces of 1µM OT-induced CA2 fEPSP responses in CD and HFD-fed mice. (L) Percentage of 1µM OT-induced fEPSP potentiation for each slice recording (measured at 10 last minutes of recording normalized to baseline) (CD: n=13,7, one-sample t-test p=2.04E-5 baseline vs end; HFD: n=13,9, Wilcoxon-rank test p=2.44E-4 baseline vs end; CD vs HFD: Mann-Whitney U test p= 0.021).

Exposure to HFD has previously been shown to alter the dendritic morphology of CA1 PNs and dentate gyrus granule cells^41^. To examine how CA2 dendritic morphology is altered following HFD-exposure, we reconstructed the dendritic arbors of CA2 PNs that had been biocytin-filled during whole-cell recordings (Fig 3B). We performed Sholl analysis and found a significant reduction in the number of intersections of basal, but not apical dendrites in HFD-exposed animals as compared to control (Fig 3C). Although the total length of apical and basal dendrites was similar in both groups, quantification of the dendritic nodes and dendritic order number revealed significant reductions for basal dendrites in the HFD-exposed mice as compared to CD (Fig 3D; CD: n=17,10, HFD: n=20,13 unpaired t-test p<0.05). These results are similar to reports of dendritic morphology changes of CA2 PNs in OTR knock-out mice^20^. Thus, we hypothesize that OT signaling in area CA2 may be altered in HFD-exposed animals.

Activation of OTRs by selective agonist application in slices has been shown to result in CA2 PN membrane depolarization and spontaneous bursting activity^24,42^ compared to CD-fed littermates (Fig 3E-H, CD: n=17,9, 7.56±0.70mV; HFD: n=17,9.χ-square test, 9, 10.56±0.86mV; Mann-Whitney U test, p=0.018). For all the CA2 PNs that had bursting activity, HFD exposure did not significantly alter burst onset, duration, number of bursts, number of APs per burst, firing rate during the bursts or the inter-burst interval (Sup Fig 8).

We then examined whether HFD exposure altered the ability of OTR activation to enhance CA2 PN intrinsic excitability. To do this, we examined the number of APs elicited with current injection as well as the intrinsic properties of CA2 PNs before and during OT application. OT application caused a leftward shift in the AP number per current injected in both CD and HFD groups (Sup Fig 9A-B). OT also induced a decrease in rheobase current, an increase in membrane resistance and an increase in sag in both CD and HFD groups (Sup Fig 9C-F). Interestingly, while OT increased AP threshold of CA2 PNs in the CD-fed mice, this did not occur in the HFD-fed mice (Sup Fig 9E). As previously reported^24^, activation of OTRs results in a change in the AP shape in CA2 PNs. In our results, AP half-width and AP decay time constants were reduced by OT application in both diet groups (Sup Fig 10A-E). Because AP half-width and decay time constants were already reduced at baseline in HFD-fed mice, these measures were also lower in HFD-fed mice compared to CD-fed mice upon OT application (Sup Fig 10D-E). This small OT-induced increase in AP decay time may potentially result in less calcium entry per individual AP, and hence, alter the link between OTR activation and intracellular signaling of CA2 PNs, the change is relatively small compared to the large increase in synaptic transmission in area CA2 following HFD exposure. Thus, while some aspects of OT-induced enhanced intrinsic excitability are altered in HFD conditions, we doubt that these alone are sufficient to account for the large effect of HFD on SRM.

### A HFD impairs OT-mediated long-term potentiation of CA3-CA2 synapses

OTR activation in slices has been shown to result in a long-term potentiation (LTP) of glutamatergic transmission at the CA3-CA2 and EC-CA2 synapses, as well as a depression of inhibitory transmission recruited by EC^20,25^. To determine if HFD-exposure impaired LTP of CA3-CA2 responses, we measured the change in synaptic transmission following application of 1 µM OT while maintaining the cells at -70 mV in current-clamp mode. In slices prepared from HFD-fed mice, application of OT resulted in reduced potentiation of CA3-CA2 synapses as compared to CD-fed mice (Fig 3I-J, 40-50min: CD: n=15,13 one-sample t-test, p=1.0E-4 different from baseline vs; HFD: n=15,12 one-sample t-test p=0.0023 different from baseline HFD: unpaired t-test p=0.0081). To better understand the synaptic mechanisms of this change, we tested whether a HFD altered OT-induced changes in inhibitory transmission by recording cells in voltage clamp mode at +10 mV to isolate IPSCs and blocked all excitatory transmission with AMPAR and NMDAR blockers (AP-5 and NBQX). OT application led to a long-term depression of IPSCs onto CA2 PNs, as had previously been reported^20^ upon stimulation of the EC input to CA2 for both HFD and CD-fed mice (Sup Fig 11 A-B, 30-40min: CD: n=7,5 one-sample t-test p=0.0049 different from baseline; HFD: n=8,6, one-sample t-test p=0.0041 different from baseline; CD vs HFD: unpaired t-test p= 0.63). These results suggest that HFD exposure does not alter the effect of OTR activation on inhibitory transmission in area CA2. We performed extracellular recordings of excitatory transmission onto area CA2 in the presence of GABA_R_ blockers and confirmed that 1µM OT application induced a smaller potentiation of transmission in HFD-fed mice as compared to the potentiation induced in their CD-fed littermates (Fig 3K-L, CD: n=13,7, one-sample t-test p=2.0E-5 different from baseline; HFD: n=13,9, Wilcoxon rank test p=2.4E-4 different from baseline; CD vs HFD: Mann-Whitney U-test p= 0.021).

We have previously demonstrated a functional link between CA2 PN AP firing and eCB-mediated long term depression of inhibition (iLTD)^34^. It may be that some increase in overall field potential that we see with OT application may be due to eCB-mediated iLTD. To answer this question, we recorded field potentials in area CA2 before and after application of 1 µM OT in the presence of the cannabinoid receptor type 1 (CB1R) inverse agonist, AM251, for both CD-fed (Sup Fig 12 A-B, CD OT: n=14,12, one-sample t-test p=0.0021 different from baseline; CD OT + AM251: n=8,8, one-sample t-test p=0.0625 different from baseline; CD OT vs CD OT + AM251: unpaired t-test p= 0.7) and HFD-fed (Sup Fig12 C-D, HFD OT: n=9,3, one-sample t-test p=0.0002 different from baseline; HFD OT + AM251: n=7,3, one-sample t-test p=0.0842 different from baseline; HFD OT vs HFD OT + AM251: unpaired t-test p= 0.6136) conditions. We observed that the change in field potential was unaffected, indicating there is no contribution of eCB-mediated iLTD to the LTP induced by OT activation alone.

Because we saw that HFD exposure resulted in increased excitatory transmission at the CA3-CA2 synapse (see Fig 1F-I), we postulated that perhaps the reduced LTP following 1 µM OT application may be due to an occlusion, or ceiling effect of the OT-dependent potentiation in HFD-fed mice. We tested whether electrical stimulation led to higher OT release in slices of HFD-fed mice compared to CD-fed mice. Antagonizing OTRs via application of 1µM of L-37 did not alter excitatory transmission onto CA2 PNs in HFD or CD-fed mice (Sup Fig 13), suggesting that no basal OT is released with electrical stimulation in hippocampal slices in area CA2 in our experiments.

### Application of 1µM OT is permissive for eCB-mediated plasticity in area CA2 in CD but not HFD-fed mice

It has previously been demonstrated that expression of OTRs in CA2/CA3a PNs is necessary for SRM^20,26,27^. Furthermore, it has been shown that synaptic plasticity of inhibitory synapses is also required for SRM^34^. To better understand the changes occurring in HFD-fed mice, we examined the two forms of iLTD in area CA2: delta-opioid receptor (DOR)-mediated and eCB-mediated long-term depression of inhibitory transmission (iLTD). These synaptic plasticities occur on separate populations of inhibitory terminals. Furthermore, the eCB-mediated iLTD in area CA2 requires the PN to fire APs, causing calcium influx and eCB release (Sup Fig 14A-B). The DOR-mediated iLTD has been shown to be permissive of the eCB-mediated plasticity, as it reduces inhibitory transmission enough to allow CA2 PNs to fire APs. We found that exposure to HFD had no significant effect on the DOR-mediated LTD at inhibitory terminals in area CA2, as examined by input stimulation (Sup Fig 14C). HFD exposure also had no effect on eCB-mediated iLTD, as tested by ACEA application (Sup fig 14D), direct depolarization of the post-synaptic CA2 PN to fire APs (Sup Fig 14E) or input stimulation following DOR iLTD induction (Sup Fig 14F). Thus, dietary conditions had no effect on inhibitory synaptic plasticities in area CA2.

Activation of OTR leads to membrane depolarization and AP firing, and this may be permissive for eCB release if paired with synaptic transmission. To do this, we used a sub-threshold tetanus induction protocol of two trains of 25 pulses at 10 Hz that did not consistently induce eCB-mediated iLTD in both CD and HFD conditions (Sup Fig 15). In CD-fed mice, we applied 1µM OT 5 minutes before applying the sub-threshold induction protocol. This induced a strong increase in fPSP amplitude at CA3-CA2 synapses (Fig 4A-C, 50-60 min: 39.23±5.38% increase, one-sample t-test p=9.6E-5 different from baseline, CD OT+tetanus: n=10,8) that was blocked by the CB1R inverse agonist AM251 (Fig 4C, 9.00±6.07% increase, one sample t-test p=0.16 different from baseline; OT+tetanus+AM251: n=8,7; unpaired t-test p=0.0022 for difference between groups). Thus, we conclude that this increase in fPSP is due to the eCB-mediated disinhibition of the CA3-CA2 synapse. We verified that the increase of fPSP amplitude was not due to the sub-threshold tetanus alone or OT application alone with interleaved control experiments. The largest increase in field potential occurred only with the combination of both OT application and subsequent sub-threshold tetanus (Fig 4D, unpaired t-test p=0.024 for OT+tetanus versus OT only; p=0.0015 for OT+tetanus versus tetanus alone). Thus, a brief application of 1µM OT is therefore permissive for eCB-mediated iLTD in area CA2.

**Figure 4.**
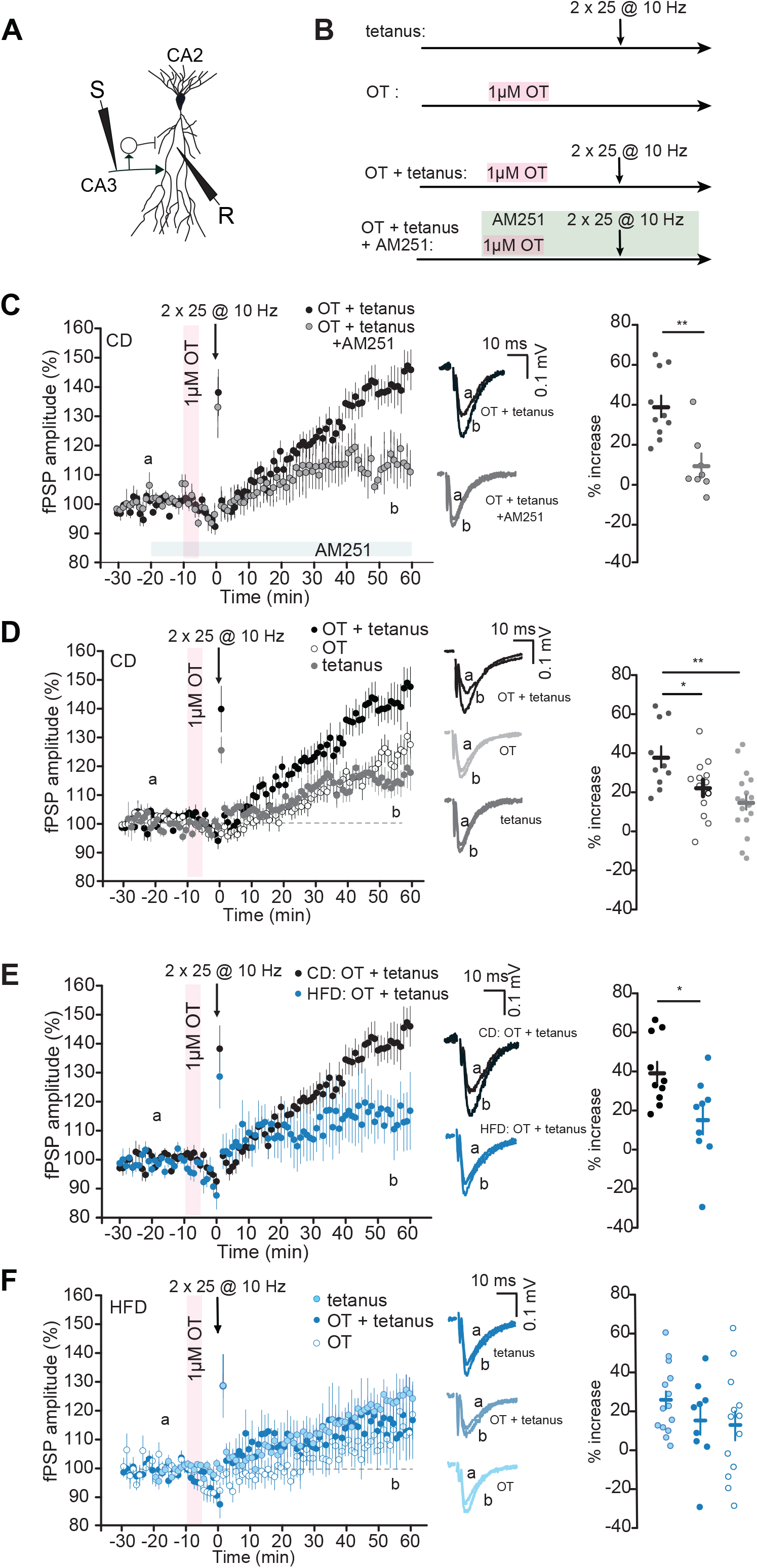
OT is permissive for eCB plasticity in CA2 of CD but not HFD-fed mice. (A) Schematic of CA2 fPSP recording with SC stimulation. (B) Experimental protocols to test for OT-induced eCB plasticity. (C) 5-minute application of 1µM OT 10 minutes before an induction protocol (2*25 @10 Hz, 20V) allows for CB1R-dependent fPSP potentiation. Middle: example traces of fPSPs before and after induction protocol with and without AM251 (CB1R antagonist). Right: 5-minute application of OT + induction allows for long-term potentiation of fPSPs (one-sample t-test p=9.6E-5 for difference with baseline). This effect is blocked if AM251 is applied (one-sample t-test p= 0.16 for OT+AM251 difference with baseline; unpaired t-test p=0.0022 for difference between groups; OT+10 Hz: n=10,8: OT+10 Hz+AM251: n=8,7). (D) OT+10 Hz protocol allows for stronger fPSP potentiation than OT alone or induction alone in CD mice. Middle: example traces of fPSPs before and after induction protocol for the three conditions. Right: 5-minute application of OT+10 Hz allows for stronger long-term potentiation of fPSPs than OT alone or 10 Hz induction alone (one-sample t-test p=7.8E-4 for OT alone difference with baseline; one-sample t-test p=0.0011 for 10 Hz alone difference with baseline; unpaired t-test p=0.024 for OT+10 Hz versus OT only; unpaired t-test p=0.0015 for OT+10 Hz versus 10 Hz alone; OT+10 Hz: n=10,8: OT only: n=16,12; 10 Hz only: n=17,16). (E) 5-minute application of 1µM OT 10 minutes before an induction protocol (2*25 @10 Hz, 20V) allows for fPSP increase in CD but not HFD-fed mice. Middle: example traces of fPSPs before and after induction protocol in CD and HFD-fed mice. Right: 5-minute application of OT + induction allows for increase of fPSPs in CD-fed mice, but not in HFD-fed mice (unpaired t-test p= 0.03 for difference between groups; one-sample t-test p=9.6 E-5 for CD OT+10Hz for difference with baseline; one-sample t-test p= 0.065 for HFD OT+10 Hz for difference with baseline; CD OT+10 Hz: n=10,8: HFD OT+10 Hz: n=9,7). (F) OT+10 Hz, OT only and 10 Hz only protocols allow for the same extent of fPSP increase in HFD mice. Middle: example traces of fPSPs before and after induction protocol for the three conditions. Right: 5-minute application of OT+10 Hz allows for similar increase of fPSPs in three conditions (one-sample t-test p= 0.13 for HFD OT alone difference with baseline; one-sample t-test p= 2.8E-4 for HFD 10 Hz alone difference with baseline; unpaired t-test p=0.8 for OT+10 Hz versus OT only; unpaired t-test p=0.2 for OT+10 Hz versus 10 Hz alone; HFD OT+10 Hz: n=9,7: OT only: n=14,10; 10 Hz only: n=15,14). All experiments were done in the presence of naltrindole.

In HFD-exposed animals, however, no eCB-mediated iLTD was observed with the sub-threshold tetanus following 1 µM OT application (Fig 4E, CD OT+tetanus: n=10,8, one-sample t-test p=9.6E-5 different from baseline; HFD OT+tetanus: n=9,7, one-sample t-test p=0.0651 different from baseline; CD OT+tetanus vs HFD OT+tetanus: unpaired t-test p=0.0308). We propose that compromised OT signaling may prevent the eCB-mediated plasticity from occurring in these conditions in HFD mice. In HFD-exposed mice, OT application followed by a subthreshold tetanus produced no additional increase in fPSP than the sub-threshold tetanus alone, or OT application alone (Fig 4F, HFD OT+tetanus: n=9,7, one-sample t-test p=0.0651 different from baseline; HFD OT: n=14,10, one-sample t-test p=0.1353 different from baseline; HFD tetanus: n=15,14, one-sample t-test p=0.00028 different from baseline; HFD OT+tetanus vs HFD OT: unpaired t-test p=0.8012; HFD OT+tetanus vs HFD tetanus: unpaired t-test p=0.2085).

### Activation of hM4D(Gi) DREADDs in area CA2 rescues OT permissive eCB-mediated iLTD in HFD-fed mice

We hypothesize that during social exploration, OT is released into area CA2, and the activation of OTRs is permissive for eCB-mediated iLTD required for SRM in CD mice. In HFD-fed mice, OTR activation is not permissive for eCB-mediated iLTD, and consistently, SRM is compromised (Fig 2). We found that activation of hM4D(Gi) DREADDs in CA2 PNs of HFD mice can rescue SRM. Using the same chemogenetic strategy, we injected an AAV vector expressing floxed-hM4D(Gi) DREADDs bilaterally into the dCA2 of both male and female CD-and HFD-fed Cacng5-Cre mice (Fig 5A). We then tested the effect of activating hM4D(Gi) DREADDs on OT-permissive eCB-mediated iLTD in acute hippocampal slices. We found that Gi-DREADD activation completely restored eCB-mediated iLTD following 1 µM OT application in HFD-fed animals (fig 5B-E, HFD CNO OT+tetanus: n=20,9, one-sample t-test p<0.0001 different from baseline; HFD OT+tetanus: n=14,9, one-sample t-test p=0.0005 different from baseline; HFD CNO OT+tetanus+AM251: n=12,9, one-sample t-test p<0.0001 different from baseline; HFD CNO OT+tetanus vs HFD OT+tetanus: unpaired t-test p=0.047; HFD CNO OT+tetanus vs HFD CNO OT+tetanus+AM251: unpaired t-test p=0.0118; HFD OT+tetanus vs HFD CNO OT+tetanus+AM251: unpaired t-test p=0.1754). The involvement of eCB plasticity was confirmed by the application of AM251 (Fig 5D-E). In these experiments, we saw no statistical difference in the plasticity changes between males and females, even though the social novelty preference was altered in a sex-dependent manner. Similar to what we have previously reported^34^, activation of Gi-DREADD in CA2 PNs prevents eCB-mediated iLTD in CD conditions, even following application of 1 µM OT (Fig 5F-G, CD CNO OT+tetanus: n=7,5, one-sample t-test p=0.0019 different from baseline; CD OT+tetanus: n=10,10, one-sample t-test p=0.0005 different from baseline; CD CNO OT+tetanus+AM251: n=6,5, one-sample t-test p=0.0143 different from baseline; CD CNO OT+tetanus vs CD OT+tetanus: unpaired t-test p=0.016; CD CNO OT+tetanus vs CD CNO OT+tetanus+AM251: unpaired t-test p=0.2903; CD OT+tetanus vs CD CNO OT+tetanus+AM251: unpaired t-test p=0.1845). These results are consistent with our behavioral data, showing that HFD-fed animals have rescued SRM and CD-fed animals have compromised SRM with activation of hM4D(Gi) DREADDs in area CA2 (fig 2), and indicate that the link between intracellular signaling and eCB release is altered in HFD-fed mice.

**Figure 5.**
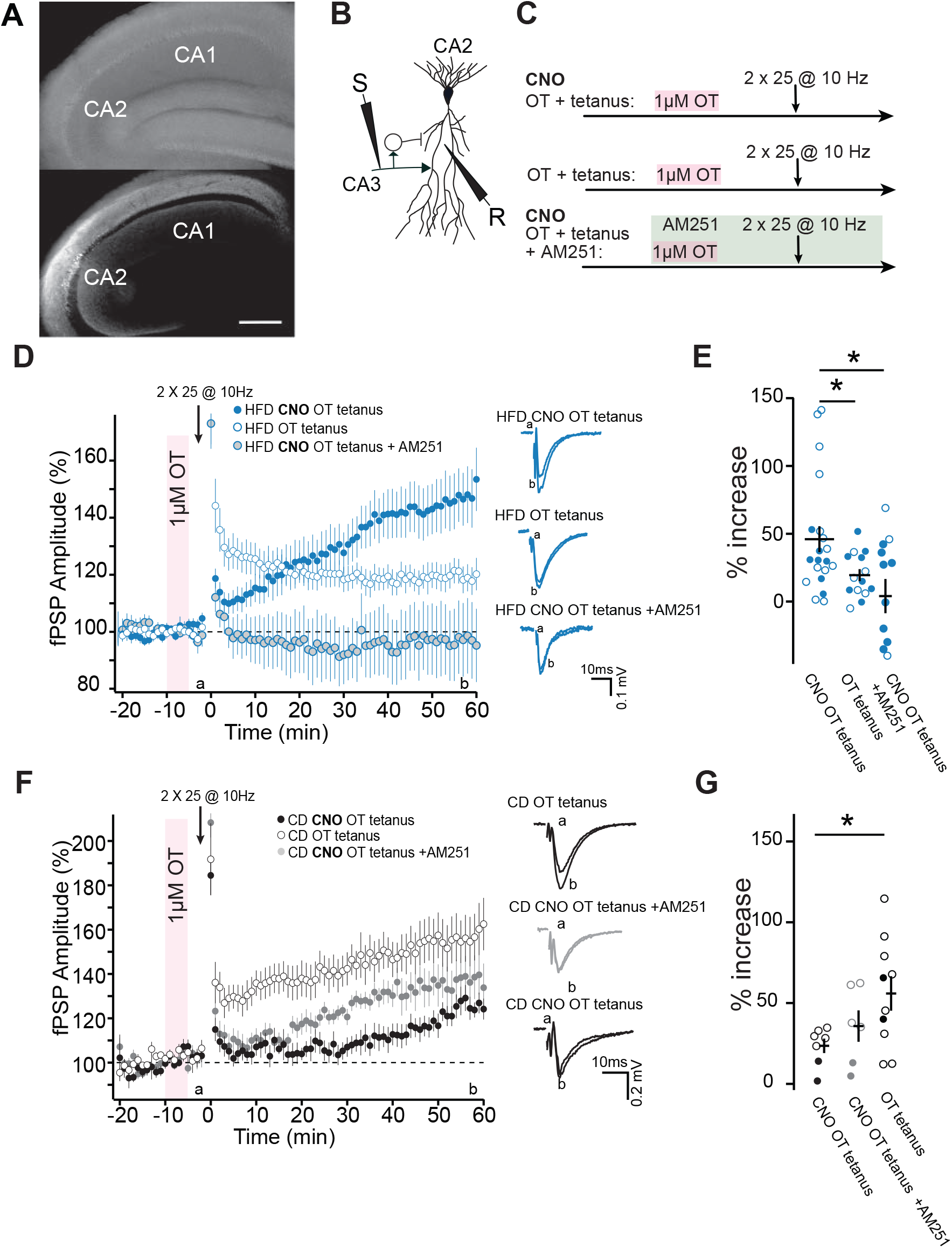
Activation of Gi-DREADD in CA2 PNs restores OT-permissive eCB-mediated plasticity in HFD-fed mice. (A) Representative image of a hippocampal slice prepared from an animal that had Gi-DREADD bilaterally injected into hippocampal area CA2 of Cacng5-cre mice. Top panel, brightfield image with area CA2 and CA1 labeled. Lower panel, fluorescent image of mCherry-Gi-DREADD selectively expressed in area CA2. Scale bar is 500 µm. (B) Schematic of CA2 fPSP recording upon SC stimulation. (C) Experimental protocols to test for activation of Gi-DREADD with or without 10 µM Clozapine-N-Oxide applied throughout the experiment, combined with 5-minute of 1 µM OT application, and with or without 4 µM AM251 application to confirm eCB-mediated plasticity. (D) Left, in slices from HFD-fed mice, activation of Gi-DREADD receptors in CA2 PNs by application of CNO restores CB1R-dependent increase in field potentials following a 5-minute application of 1µM OT 10 minutes before a sub-threshold stimulation (2*25 @10 Hz, 20V). This effect is prevented by blocking CB1Rs with AM251. Right: example traces of fPSPs for all three conditions. (E) Percentage of increase from baseline for all three conditions (HFD CNO OT and tetanus: n=20,9, 146.0±9.535, one sample t-test p<0.0001; HFD OT with tetanus n=14,9, 119.4±4.415 one-sample t-test p=0.0005, HFD CNO with OT, tetanus and AM251: n=12,9, 103.03±12.65 one-sample t-test p<0.0001; for difference between groups; HFD CNO OT and tetanus vs HFD OT and tetanus: unpaired t-test p=0.047; HFD CNO OT and tetanus vs HFD CNO with OT, tetanus and AM251: unpaired t-test p=0.0118; HFD OT and tetanus vs HFD CNO with OT, tetanus and AM251: unpaired t-test p=0.1754). (F) Left, in slices from CD-fed mice, activation of Gi-DREADD receptors in CA2 PNs by application of CNO prevents CB1R-dependent increase in field potentials following a 5-minute application of 1µM OT 10 minutes before a sub-threshold stimulation (2*25 @10 Hz, 20V). In CD-fed mice, the activation of Gi-DREADDs prevents eCB-mediated plasticity. Right: example traces of fPSPs for all three conditions. (G) Percentage of increase from baseline for all three conditions (CD CNO OT and tetanus: n=7,5, 123.51±4.45, one sample t-test p=0.0019; CD OT with tetanus n=10,10, 155.91±10.67 one-sample t-test p=0.0005, CD CNO with OT, tetanus and AM251: n=6,5, 135.73±9.715 one-sample t-test p=0.0143; for difference between groups; CD CNO OT and tetanus vs CD OT and tetanus: unpaired t-test p=0.0161; CD CNO OT and tetanus vs CD CNO with OT, tetanus and AM251: unpaired t-test p=0.2903; CD OT and tetanus vs CD CNO with OT, tetanus and AM251: unpaired t-test p=0.1845).

We did not see any sex-dependent differences for the permissive effect of OT of eCB-mediated plasticity (Two-way ANOVA, sex-effect, F _(1, 20)_=2.353, p=0.1407). Nor did we see any sex-dependent difference in the effect of Gi-DREADD activation (Two-way ANOVA, sex-effect, F _(1,23)_=1.333, p=0.2602). This is consistent with the behavior data in that both male and female mice have compromised SRM, and were rescued with Gi-DREADD activation, even though the social novelty preference was altered in females.

### 10 µM OT can rescue eCB-mediated plasticity in HFD-fed mice

Our results support the idea that OT signaling is altered in HFD-fed mice, as indicated by the reduced ability of 1 µM OT to increase synaptic transmission, as well as the absence of eCB-mediated plasticity following 1 µM of OT application. We examined whether a higher concentration of OT could rescue the OT-induced eCB plasticity in HFD-fed mice. To test this, we used the same experimental setup, except with a 10 µM OT application preceding the sub-threshold tetanus induction protocol (Fig 6A-B). In CD conditions, we found that the resulting increase in field potential with both tetanus and 10 µM OT was the same as what we observed with application of 10 µM OT alone (Fig 6C; CD OT+tetanus: n=7,5, one-sample t-test p=0.0003 different from baseline; CD OT: n=10,7, one-sample t-test p=0.0004 different from baseline; CD tetanus: n=7,5, one-sample t-test p=0.1641 different from baseline; CD OT+tetanus vs CD OT: unpaired t-test p=0.9335; CD OT vs CD+tetanus: unpaired t-test p=0.0236; CD OT+tetanus vs CD tetanus: unpaired t-test p=0.0189).

**Figure 6.**
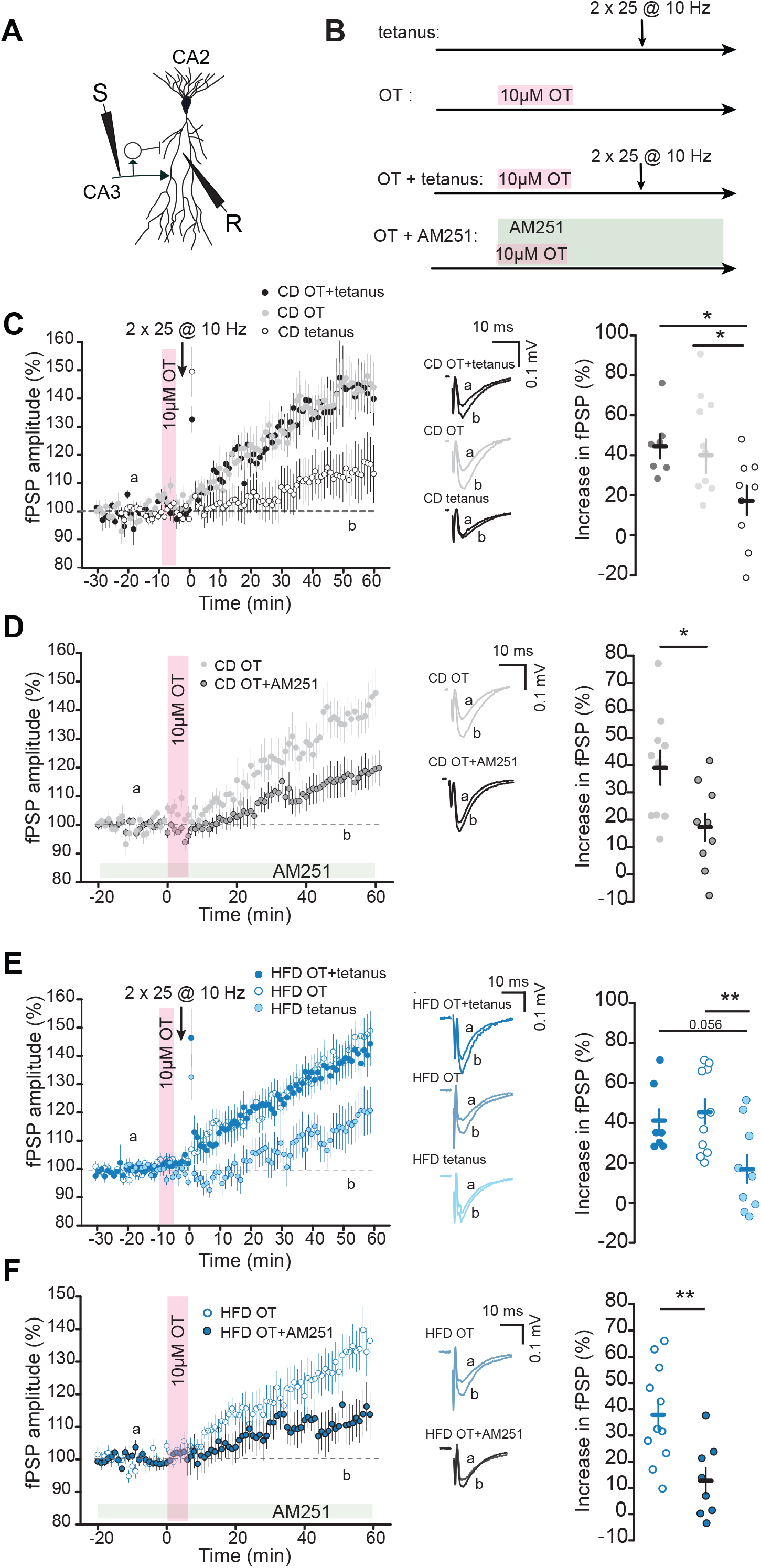
10 µM concentration of OT allows for eCB-mediated plasticity in CA2 in CD and HFD-fed mice. (A) Schematic of CA2 fPSP recording upon SC stimulation. (B) Experimental protocols to test for OT-induced eCB plasticity. (C) 5-minute application of 10µM OT 10 minutes before an induction protocol (2*25 @10 Hz, 20V) elicits long-term increase in the fPSP to the same extent as 10µM OT application alone in CD-fed mice. 10 Hz induction alone does not result in an increase in the amplitude of fPSPs (44.60±5.99%, one-sample t-test p= 3.4E-4 for OT+10 Hz baseline vs 50-60min; 45.52±7.93%, one-sample t-test p=4.0E-4 for OT alone baseline vs 50-60min; 14.63±9.28%, one-sample t-test p= 0.16 for 10 Hz alone baseline vs 50-60min). Middle: example traces of fPSPs before and after induction protocol. Right: 5-minute application of 10µM OT + 10 Hz and of 10µM OT alone allows for larger increase in potential than 10 Hz alone (unpaired t-test p= 0.023 for OT alone vs 10 Hz alone; p=0.018 for OT+10 Hz vs 10 Hz alone; p=0.93 for OT alone vs OT+10 Hz; OT+10 Hz: n=7,5: OT alone: n=10,7; 10 Hz alone: n=7,5). (D) Left: AM251 hinders 10µM OT-induced increase in the field potential by preventing eCB iLTD in CD-fed mice (50-60min; OT only: 39.06±6.26%; one-sample t-test p= 2.4E-4 different from baseline; OT+AM251: 17.37±5.31%; one-sample t-test p=0.010 different from baseline; OT only: n=10,7; OT+AM251: n=9,6). Middle: example traces before and after OT application with and without AM251 application. Right: Percentage of 10µM-OT induced fPSP potentiation is reduced when blocking CB1Rs (unpaired t-test p=0.018 OT only vs OT+AM251). (E) 5-minute application of 10µM OT 10 minutes before an induction protocol (2*25 @10 Hz, 20V) elicits long-term increase in the field potential that is the same magnitude as 10µM OT application alone in HFD-fed mice. 10 Hz induction alone elicits a smaller increase in potential of fPSPs (50-60min: 41.17±6.51%, Wilcoxon-rank test p=0.022 for HFD OT+10 Hz different from baseline; 45.82±6.22%, one-sample t-test p=2.4E-5 for HFD OT alone different from baseline; 16.94±7.35%, one-sample t-test p= 0.032 for HFD 10 Hz alone different from baseline). Middle: example traces of fPSPs before and after induction protocol. Right: 5-minute application of 10µM OT + 10 Hz and of 10µM OT alone allows for stronger increase in field potential than 10 Hz alone in HFD-fed mice (two-sample t-test p= 0.0074 for HFD OT alone vs HFD 10 Hz alone; Mann-Whitney U-test p=0.056 for HFD OT+10 Hz vs HFD 10 Hz alone; Mann-Whitney U-test p=0.78 for HFD OT alone vs HFD OT+10 Hz; HFD OT+10 Hz: n=7,6: HFD OT alone: n=11,7; HFD 10 Hz alone: n=9,7). (F) Left: AM251 hinders 10µM OT-induced increase of CA2 fPSP amplitude in HFD-fed mice (50-60min: HFD OT only: 37.96±5.62%; one-sample t-test p= 4.8E-5 different from baseline; HFD OT+AM251: 12.72±4.99%; one-sample t-test p=0.043 different from baseline; HFD OT only: n=11,7; HFD OT+AM251: n=8,4). Middle: example traces before and after OT application with and without AM251 application. Right: % 10µM-OT induced fPSP potentiation is reduced when blocking CB1Rs in HFD-fed mice (unpaired t-test p=0.0051 HFD OT only vs HFD OT+AM251). All experiments were done in the presence of Naltrindole.

Because a higher concentration of OT elicits strong CA2 PN bursts on its own, we tested whether the 10 µM OT application alone was sufficient to induce eCB-mediated plasticity in area CA2. We found that application of the CB1R inverse agonist AM251 impaired 10 µM OT-induced increase of CA3-CA2 fPSP amplitude (Fig 6D; CD OT: n=10,7, one-sample t-test p=0.0002 different from baseline; CD OT+AM251: n=9,6, one-sample t-test p=0.010 different from baseline; CD OT vs CD OT+AM251: unpaired t-test p=0.0183). We next examined how the coupling between 10µM OT and eCB-mediated plasticity was altered by HFD-exposure. In HFD-fed mice, application of 10 µM OT led to the same increase in fPSP amplitude with or without the sub-threshold tetanus (Fig 6E; HFD OT+tetanus: n=7,6, Wilcoxon test p=0.022 different from baseline; HFD OT: n=11,7, one-sample t-test p=4E-4 different from baseline; HFD tetanus: n=9,7, one-sample t-test p=0.0328 different from baseline; HFD OT+tetanus vs HFD OT: Mann-Whitney U-test p=0.7858; HFD OT vs HFD tetanus: unpaired t-test p=0.0074; HFD OT+tetanus vs HFD tetanus: Mann-Whitney U-test p=0.0567) and was sufficient to induce eCB plasticity in HFD-fed mice (Fig 6F, HFD OT: n=11,7, one-sample t-test p=4E-4: HFD OT+AM251: n=8,4, one-sample t-test p=0.0439; HFD OT only vs HFD OT+AM251: unpaired t-test p=0.0051). Thus, these results indicate that both the OT-mediated LTP and the e-CB-mediated disinhibition are both potentially dysfunctional in HFD-fed mice and can be rescued with higher concentration OT application.

### Injection of a high, but not low, concentration of OT in dCA2 rescues SRM deficits in HFD-fed mice

We have shown that in HFD-exposed mice, 10 µM OT is able to rescue in dCA2 both the OT-mediated potentiation as well as enable eCB-mediated disinhibitory increase in fPSPs. This is in contrast to 1 µM OT application. We hypothesized that an increased OT during social interaction may rescue the SRM deficits of HFD-fed mice. To test this, we bilaterally implanted cannulas in dCA2 (Fig 7A-B) and delivered either vehicle, 1 ng or 10 ng OT (corresponding to 5 µM and 50 µM OT, respectively) in 0.2 µL bilaterally 10 minutes prior to the training phase of the SRM test in both CD-fed and HFD-fed mice. Injection of vehicle or OT at both concentrations did not alter sociability either in the CD-fed or HFD-fed mice (Fig 7C-D; CD vehicle: n=7, one-sample t-test p=1.00E-04; CD OT 5μM: n=7, one-sample t-test p=4.00E-04; CD OT 50μM: n=8, one-sample t-test p=2.00E-04; HFD vehicle: n=7, one-sample t-test p=1.00E-04; HFD OT 5μM: n=6, one-sample t-test p=8.90E-03; HFD OT 50μM: n=9, one-sample t-test p=0.0001; Two-way ANOVA F _(2,38)_= 1.51 p=0.23). During the SRM testing, 3 hours later, we observed that vehicle-infused CD-fed mice exhibited normal ability to discriminate between familiar and novel conspecifics. However, the CD-fed animals that received either the 5 µM or 50 µM OT injection displayed impaired SRM, as evidenced by their poor discrimination between the familiar and novel stimuli. In HFD-fed mice, those injected with vehicle displayed impaired SRM. Likewise, SRM performance was still impaired with 5 µM OT injection. In contrast, 50 µM OT injection in dCA2 rescued SRM deficits in HFD-fed animals (Fig 7E-F CD vehicle: n=7, one-sample t-test p=3.50E-02; CD OT 5μM: n=7, one-sample t-test p=1.60E-01; CD OT 50μM: n=8, one-sample t-test p=3.00E-01; HFD vehicle: n=7, one-sample t-test p=4.90E-01; HFD OT 5μM: n=6, one-sample t-test p=2.90E-01; HFD OT 50μM: n=9, one-sample t-test p=0.0023; Two-way ANOVA F _(2,38)_=4.59 p=0.0164).These results indicate that exogenous OT injection in dCA2 in CD-fed animals during social interaction can effectively perturb SRM. However, in the HFD-fed mice, the elevated OT is sufficient to rescue SRM deficits.

**Figure 7.**
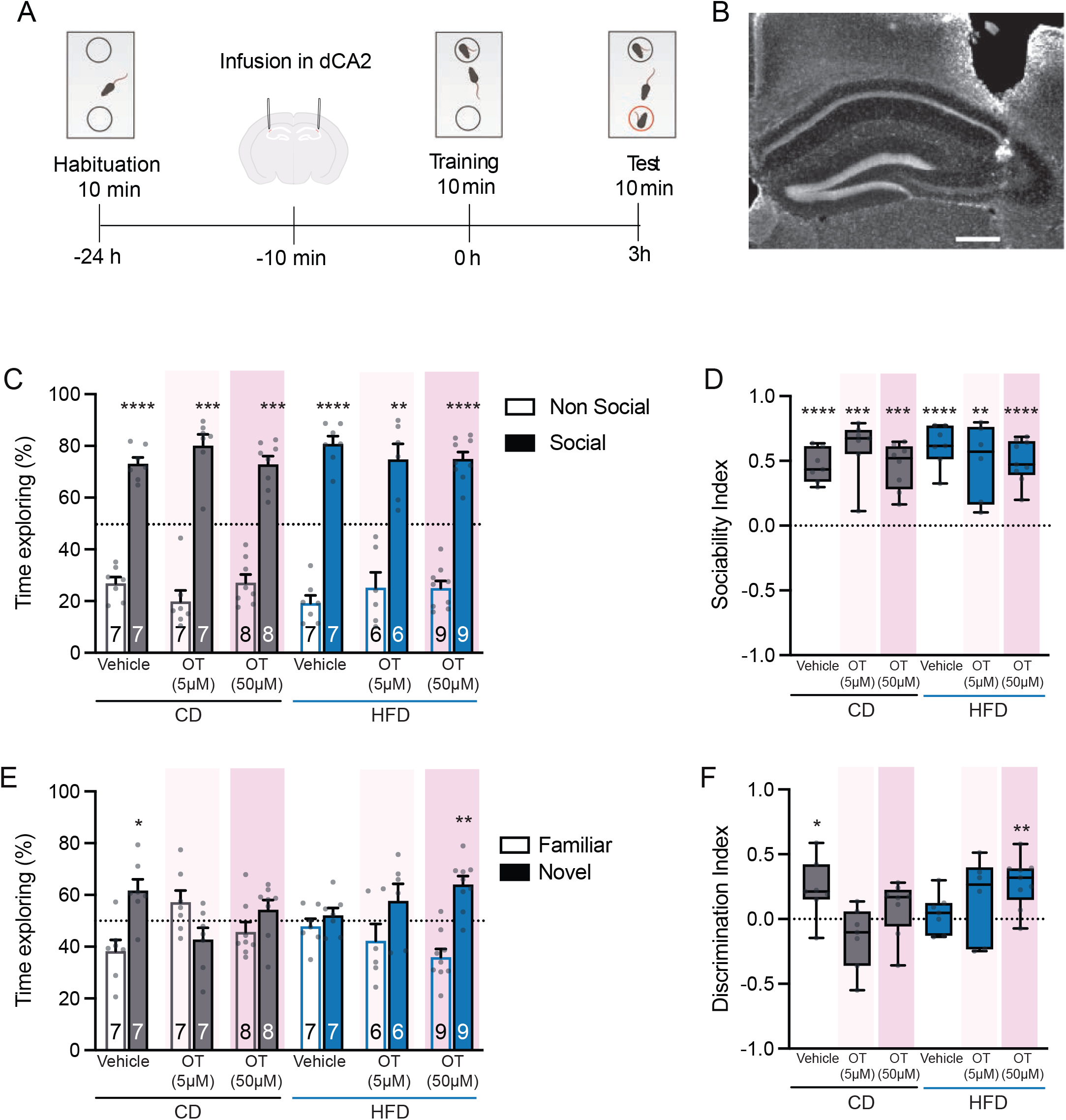
High, but not low, concentration of OT injected bilaterally in dCA2 rescues SRM in HFD-fed mice, and both low and high OT injection in dCA2 impairs SRM in CD-fed mice. (A) Behavioral paradigm. (B) Example image of cannula implantations in dCA2. Scale bar corresponds to 100 µm. (C) HFD and CD mice, injected either vehicle, low or high concentration OT, explore the social stimulus a greater amount of time compared to the non-social one for comparison between time exploring social and non-social stimulus in all groups. (D) Sociability index for social versus non-social stimuli is different than 0 in both CD and HFD-fed mice (CD vehicle: n=7, one-sample t-test p=1.00E-4; CD OT 5μM: n=7, one-sample t-test p=4.00E-4; CD OT 50μM: n=8, one-sample t-test p=2.00E-4; HFD vehicle: n=7, one-sample t-test p=1.00E-4; HFD OT 5μM: n=6, one-sample t-test p=8.90E-3; HFD OT 50μM: n=9, one-sample t-test p=0.0001; Two-way ANOVA F _(2,38)_= 1.51 p=0.23). (E) In the test phase, CD vehicle mice explore the novel mouse a greater amount of time compared to the familiar one. CD-low OT and CD-high OT mice explore the novel and familiar mice the same amount of time. HFD-vehicle mice and HFD-low OT mice do not discriminate between the novel and the familiar mouse. HFD high-OT mice explore the novel mouse over the familiar mouse. (F) Discrimination index is higher than 0 in CD-vehicle and HFD-high OT, but not in CD-low OT, CD-high OT or HFD-vehicle and HFD-low OT mice (CD vehicle: n=7, one-sample t-test p=3.50E-2; CD OT 5μM: n=7, one-sample t-test p=1.60E-1; CD OT 50μM: n=8, one-sample t-test p=3.00E-1; HFD vehicle: n=7, one-sample t-test p=4.90E-1; HFD OT 5μM: n=6, one-sample t-test p=2.90E-1; HFD OT 50μM: n=9, one-sample t-test p=0.0023; Two-way ANOVA F _(2,38)_=4.59 p=0.0164). The number of animals used per group is shown on the figure.

## DISCUSSION

### Cellular mechanism linking oxytocin, eCB plasticity and social recognition memory

Our results reveal a link between OT neuromodulation and synaptic plasticity in area CA2. It is known that both functional OTRs and CB1Rs in area CA2 are necessary for social memory formation^20,26,27,34^. Here, we demonstrate that OTR activation in area CA2 is permissive for eCB-mediated iLTD at GABAergic synapses. This OT-induced eCB plasticity is likely highly relevant because OT signaling has critical importance for social behaviors. In the ventral striatum, another area receiving OT projections from the PVN, stimulation of OT release has also been shown to drive eCB-mediated signaling^43^. Here, we demonstrated that a combination of 1 µM OT application prior to a sub-threshold induction protocol resulted in eCB iLTD (see schema illustrating the results and potential mechanism; Sup Fig 16). We demonstrate that this concentration of OT is sufficient to depolarize CA2 PNs and to induce burst AP firing. This depolarization of CA2 PNs and AP firing are likely allowing for the sufficient intracellular calcium entry with concomitant synaptic transmission to enable eCB release by CA2 PNs^34^. We propose that OT is released in dCA2 during social interactions and enables eCB-mediated iLTD, allowing for encoding of social information.

### dCA2 hyperexcitability impairs SRM in HFD-fed mice

While many attributes of CA2 PNs do not seem to be altered in HFD-fed mice, stimulation of afferents from the EC and dorsal CA3 are aberrantly large. Changes in the paired-pulse ratio indicate potential changes in pre-synaptic neurotransmitter release. Both of these inputs onto area CA2 have been reported to affect SRM^23,44^. Here we showed that expression of Gi-DREADDs during memory formation of a social encounter impaired SRM in CD-fed mice. In HFD-fed mice, Gi-DREADD expression in CA2 PNs restored the ability of mice to discriminate between a novel and familiar mouse. Whereas CA2 inhibition in HFD-fed males restored social novelty preference during SRM test, similar manipulation in HFD-fed females is associated with social familiarity preference. Interestingly, the activation of Gi-DREADDs was able to allow 1µM OT application to be permissive for eCB-mediated plasticity in slices prepared from both HFD-fed males and females. For the CD-fed animals, Gi-DREADD activation was detrimental to both OT-permissive eCB plasticity and SRM. While this result goes hand-in-hand with previously published reports showing that not only is CA2 activity necessary for SRM, but that its activity must be finely tuned in order to engage in proper social memory formation^44^, the sex effect on social novelty/familiarity preference in HFD-fed mice remains to be more thoroughly investigated.

### Two distinct CA2 strategies to rescue HFD-induced SRM deficits

It seems surprising that both chemogenetic inhibition of CA2 PNs and injections of a high dose of OT in area CA2 rescue SRM impairments in HFD-fed mice, as OT application increases excitability and spontaneous burst firing of CA2 PNs^24^. This apparent contradiction may be related to an insufficient signal-to-noise ratio problem. In HFD-fed mice, area CA2 exhibits aberrant basal excitatory transmission. This “noise” may be too high for social-stimulus mediated signals, such as “low” (1 µM) OT signaling, to have a sufficiently strong effect on CA2 firing upon social interactions (Sup Fig 16). When CA2 PNs are silent, as they are in CD-fed mice, lower OT signals are permissive for synaptic inputs to result in eCB plasticity. It is possible that 10 µM OT injected in CA2 rescues HFD-induced SRM deficits as it allows for the OT social signal to be sufficiently strong to overcome the background hyperexcitable noise in area CA2 and induce plasticity necessary for encoding of social information. We demonstrate how activation of Gi-DREADD in CA2 PNs of HFD-fed animals restores the ability of 1 µM OT to enable eCB-mediated plasticity. We fully acknowledge, however, that the exact cellular process altered by the HFD exposure that disrupts normal plasticity in CA2 PNs is unknown. Furthermore, which downstream signaling pathway is underlying the chemogenetic rescue is also an open question.

### Exogenous OT injection in dCA2 impairs SRM in CD-fed mice

We observed that both low (5 µM) and high (50 µM) OT infusion during the training phase of the SRM task impaired performance in CD-fed mice. Previous reports indicated that high levels of OT can trigger social amnesia^45–47^, revealing that an optimal OT level and perhaps precise timing of OTR activation during social exploration is required for SRM. Indeed, it has been demonstrated that chemogenetic activation of OT afferents in dCA2 during the training phase enhanced SRM^27^. The exogenous nature and artificially high concentration of OT application could perhaps explain these contradicting findings.

Thus, exogenous OT can have opposite behavioral consequences depending on the baseline state of CA2. In CD-fed mice, where OT signaling is functional, additional OT may exceed optimal ranges and impair SRM. In HFD-fed mice, where OT-dependent plasticity is impaired, stronger OT activation may restore sufficient signaling to permit eCB-mediated plasticity.

### A link between hyperexcitability and OT signaling impairments in HFD-fed mice

The lack of OT-induced plasticity of the CA3-CA2 synapse in HFD-fed mice may decrease CA2 excitability in some contexts: this dysregulation might have led to a homeostatic compensatory increase in excitatory hippocampal inputs onto CA2. Indeed, it seems that the hyperexcitability phenotype observed in HFD-fed mice takes a long time to develop (as 10 days or 5 weeks of a HFD does not entail larger transmission at the CA3-CA2 synapse) which could be the result of a delayed adaptation. Nevertheless, even if the hyperexcitability of area CA2 is an adaptation to a lack of OT-induced effect, it seems to be a maladaptive one as it perturbs effective encoding and adds noise to hippocampal information transfer.

### Are there OT release deficits in HFD-fed mice in vivo?

As OT release in the PVN has been reported to be reduced as a consequence of a HFD^29,48^, it is possible that PVN OT release in dCA2 may be altered as well. Furthermore, the enhanced depolarization of CA2 PNs upon OT application in HFD conditions could be a resulting compensation due to lower OT signaling. However, our data cannot distinguish between impaired OT release, altered OTR expression or disrupted downstream receptor signaling. Direct monitoring of OT dynamics in dCA2 during social exploration, may resolve this question. Alternatively, it has been demonstrated that the supramammillary nucleus (SuM) expresses OTRs, and that the CA2-projecting cells in the SuM are playing an important role in SRM^21,49^. Perhaps under HFD conditions, CA2 PN activity is elevated to such an extent that the modulatory input from the SuM is ineffective. Thus, reducing or altering CA2 PNs with Gi-DREADD activation may restore the local circuit function. Further work is required, given that the slice experiments in this study were conducted under conditions that precluded any contribution of the SuM. Thus, it may be that the injection of OT may be rescuing SRM by acting on the pre-synaptic SuM inputs. Furthermore, HFD-induced astrocyte signaling may be playing a role in this phenomenon as it has been demonstrated that astrocytes, which are important regulators of excitatory transmission, express OTRs^50–52^, and are dysregulated with metabolic stress^53,54^.

### Adolescence, high fat diet and social cognition

It has been demonstrated that hippocampal area CA2 undergoes changes in cellular composition, synaptic plasticity and intrinsic excitability during adolescence^55–57^. In this work we show that prolonged exposure to a HFD diet starting at weaning, and therefore encompassing the adolescent period, results in loss of SRM in both males and females. We postulate that the timing of this dietary stress is critical, and may be disrupting the proper development of CA2 local circuitry during this critical time. We observe that the expression of Gi-DREADDs in HFD-fed mice has a different behavioral outcome for male and female mice. Whereas CA2 inhibition in HFD-fed males rescued SRM, with the expected social novelty preference, HFD-fed female with Gi-DREADD activation in CA2 PNs display a preference for a familiar conspecific. These results should be considered in conjunction with the CA2-dependent preference for familiar conspecifics (familiar or novel mothers) in early postnatal developmental periods^58^. Furthermore, the length of HFD exposure was also shown to be important, as shorter times that did not cause weight gain also did not result in changes to CA2 properties. To our knowledge, there is little to no information about changes in CA2 physiology with adolescent HFD-exposure, and more work is required to better understand which aspects of development are altered, and the consequences these would have on OT and eCB signaling.

### Limitations of the study

One major limitation of this study is that we fully do not understand how expression of Gi-DREADDs in HFD-fed mice is able to result in restored social discrimination. While we directly demonstrate that chemogenetic inhibition restores the permissive effect of OT on eCB-mediated synaptic plasticity, our interpretation may be an over-simplification. Activation of Gi-DREADDs not only alters intrinsic excitability of neurons, but also abolishes synaptic vesicle release^59^. Thus, if transmission from CA2 PNs to downstream areas including CA1, CA3 and the lateral septum is expected to be reduced, it is unclear how eCB-mediately plasticity will have any effect at all. How much this directly applies to HFD-conditions in unclear, as Gi-DREADD expression in hippocampal PNs has been shown to rescue contextual and object memory deficits^60–63^. Furthermore, activation of Gi-DREADDs has been shown to normalize fos overactivation in HFD-fed mice^61^, indicating that chemogenetics may decrease but not completely silence neuronal activity under these conditions. Thus, many unanswered mechanistic questions remain.

## RESOURCE AVAILABILITY

### Lead Contact

Requests for further information and resources should be directed to and will be fulfilled by the lead contact, Rebecca Piskorowski.

### Materials availability

All data are available upon request to the lead contact.

## Supporting information

Total Supplemental

## ACKNOWLEDGMENTS

We dedicate this work to the memory of Vivien Chevaleyre, whose contributions to this project were invaluable. The authors thank the IBPS CEFI facility, the animal facility team of NutriNeuro lab, and specifically Gregory Artaxet, for animal care. We also want to thank Ajdia Ait-Kaci and Jullie Cognet for essential technical assistance. This work was supported by the Fondation pour la Recherche Médicale EQU202003010457 to VC and RAP and French National Research Agency (ANR-18-CE370020-01, ANR-18-CE16-0006 and ANR-21-CE16-0021-03 to VC and RAP and ANR-23-CE14-0004 HIPPOBESE to GF, VC and RAP). M Muller was the recipient of a PhD fellowship from the French Ministry of Research and Higher Education (2020-2023). AF was the recipient of a PhD fellowship (2020-2023) from the Bordeaux NeuroCampus Graduate Program, managed by the French National Research Agency (ANR-17-EURE-0028), and an extension grant from the French government in the framework of the University of Bordeaux’s France 2030 program / GPR BRAIN_2030 (2023-2024).

## AUTHOR CONTRIBUTIONS

Conceptualization: VC, GF, RAP, AF, ED, and M Moschetta. Investigation: M Muller, AF, ED, AR, VC, M Moschetta, HE, MR, PAR, and RAP. Analysis: M Moschetta, M Muller, AF, ED, VC, RAP. Writing: RAP with comments from all other authors; funding acquisition, RAP, VC and GF; resources, RAP, VC and GF; supervision, RAP, VC, GF and M Moschetta.

## SUPPLEMENTAL INFORMATION

Document S1. Supplemental Figures S1–S16.

## STAR METHODS

### EXPERIMENTAL MODEL DETAILS

All experiments were performed on 18-20 week-old Swiss (Janvier labs), Swiss Cacng5cre and C57Bl6J male and female mice. Male and female mice were housed in single-sex groups of two to five per cage where they had ad libitum access to food and water. The animal facility was temperature controlled (22°C) and maintained under a 12h light/dark cycle. All procedures were performed in accordance with institutional regulations (French Ministry of Research decision #12406– 2016040417305913 v10). At weaning (3-weeks of age), litters were divided up by sex and into two-groups: one given a control diet and one a high-fat diet (HFD) in order to have age-matched, sex-matched sibling controls. The diets were given ad libitum for 12 weeks (and in some experiments, for 10 days or 5 weeks only) until the mice were used for experiments. The HFD (D12451, Research Diet) consisted of 4.7 kcal/g. The control diet consisted of 2.9 kcal/g (A04, SAFE).

C57B6J male mice (Charles River, France) were used for behavioral experiments. They were housed in groups of five since weaning at P21 in standard breeding cages with food and water ad libitum and were placed at a constant temperature (23 ± 1°C) under diurnal conditions (light-dark: 8:00AM– 8:00PM). Mice were tested during the first half of the light period.

## METHOD DETAILS

### Hippocampal slice preparation

Animals were euthanized in accordance with institutional regulations under ketamine (20 mg/kg), xylazine (1.4 mg/kg) and isoflurane, and perfused transcardially with a modified sucrose cutting solution containing the following (in mM): Sucrose 110, NaCl 10, KCl 2.5, NaH_2_PO_4_ 1.25, NaHCO_3_ 30, HEPES 20, glucose 25, thiourea 2, Na-ascorbate 5, Na-Pyruvate 3, CaCl_2_ 0.5, MgCl_2_ 10. The hippocampi were rapidly harvested, placed in an Agar mold and cut into transverse 400 μm slices with a vibratome (Leica VT1200S, Germany) in the same sucrose cutting solution at 4°C. The slices were then transferred to an immersed-type chamber containing artificial cerebrospinal fluid (ACSF) at 32°C (in mM: 125 NaCl, 2.5 KCl, 10 glucose, 26 NaHCO_3_, 1.25 NaH_2_PO_4_, 2 Na Pyruvate, 2 CaCl_2_ and 1 MgCl_2_) for approximately 15 minutes and then maintained at room temperature for at least 1.5 hours before recording. Cutting and recording solutions were both saturated with 95% O_2_ and 5% CO_2_.

### Electrophysiological recordings

All electrophysiological experiments were performed in a recording chamber allowing perfusion with ACSF at 3 mL/min at 30°C. Data were obtained using a Multiclamp 700B amplifier and digitized using a Digidata 1550. Data were sampled at 10 kHz. pClamp10 and Axograph software were used for data acquisition.

Extracellular field recordings of postsynaptic potentials (fPSPs) were recorded in current-clamp mode using a recording patch pipette (3-5 MΩ) containing 1 M NaCl and positioned in the middle of stratum radiatum in CA2. For measure of population spike (PS) activity, the recording electrode was placed in stratum pyramidale of CA2. In order to stimulate the Schaffer collateral (SC) pathway, a monopolar stimulating electrode (containing aCSF, and with a broken-off tip) was placed in stratum radiatum of CA1 in order to antidromically excite CA3-CA2 synapses. Synaptic potentials were evoked with a constant voltage stimulating unit (Digitimer Ltd.) set at 0.1 ms at a voltage range of 10–30 V.

Plasticity was induced with high-frequency stimulation (HFS; 100 pulses at 100 Hz repeated twice) in the case of delta opioid receptors-dependent plasticity induction, or 10 Hz stimulation (100 or 25 pulses at 10 Hz repeated twice, at 30 or 20V) in the case of eCB plasticity induction. When studying eCB plasticity without pre-incubation in DPDPE, an antagonist of delta-opioid receptors (Naltrindole) was bath-perfused during the induction in order to prevent delta opioid receptors-dependent plasticity.

All whole-cell recordings of CA2 pyramidal neurons (PNs) were performed “blind.” Area CA2 has extremely dense extracellular matrix, making visually guided recordings very challenging. A recording pipette with positive pressure was inserted deeply into the pyramidal cell layer parallel to stratum so that the tip of the recording pipette was even with the end of the mossy fibers. The pipette was then stepped deeper into the slice while a voltage step was delivered through the pipette. The amplitude of a 5-10 mV current step was monitored and used to determine when the pipette was close to a neuron, positive pressure was released, and a giga-ohm seal was formed.

Whole-cell recordings were performed with either a potassium or cesium-based intracellular solution depending on the experiment. The cesium-based solution contained the following (in mM): Cs-methyl sulfonate 135, KCl 5, EGTA-KOH 0.1, HEPES 10, NaCl 2, MgATP 5, Na_2_GTP 0.4, Na_2_-phosphocreatine 10 and had a pH of 7.2 and an osmolarity of 286–295 mOsm. The potassium based one contained (in mM): 135 potassium methyl-sulfate, 5 KCl, 0.1 EGTA-Na, 10 HEPES, 2 NaCl, 5 ATP, 0.4 GTP, Na_2_-phosphocreatine 10 and had a pH of 7.2 and an osmolarity of 286–295 mOsm. The liquid junction potential was not corrected. The series resistance (typically 9-25 MΩ) was monitored throughout each experiment; cells showing a series resistance >25 MΩ or a change in series resistance > 15% were excluded from the analysis.

Post-synaptic potentials (PSPs) were recorded using K^+^-internal solution in whole-cell current clamp mode with current injected to maintain V_REST_ at -70mV. Excitatory-PSPs (EPSPs) were obtained in the same configuration in the presence of GABA_A_ and GABA_B_ blockers. Inhibitory post-synaptic currents (IPSCs) were obtained in voltage-clamp mode at +10mV (reversal potential for glutamatergic transmission) using a Cs-internal solution. In order to obtain IPSCs free of feedforward inhibition, AMPAR and NMDAR blockers were applied in the bath.

Synaptic potentials and currents were evoked by stimulation with a patch pipette filled with aCSF and positioned in the middle of CA1 stratum radiatum or stratum lacunosum moleculare to stimulate SC or the perforant path, respectively. Like for field recordings, stimulation was evoked through a constant voltage stimulating unit (Digitimer Ltd.) set at 0.1 ms at a voltage range of 10–30 V.

In order to allow for eCB plasticity under whole-cell configuration which does not allow for CA2 PN action potential firing using electrical stimulation (due to feedforward inhibition^1^), short pulses of 1200pA at 100ms intervals were injected into the cell in voltage-clamp mode in order for the CA2 PN to fire. This was done via two trains of 25 7ms-long current injections. PSP and IPSC amplitudes were normalized to the baseline amplitude in plasticity experiments. The extent of long-term plasticity was estimated by using the mean response of each cell at 40-50 min after the induction protocol for whole-cell recordings, and at 50-60 min after the induction protocol for extracellular recordings and normalizing it to mean baseline response (10 minutes before induction).

Because recordings were performed blind, biocytin (HelloBio, 4 mg/mL) was added to internal solutions in order to confirm the identity of the recorded cell post-hoc via its position as well as its morphology (see section histology). In cells recorded from with K^+^-based internal solution, electrophysiological properties of CA2 PNs were also used to confirm the identity of the cell (input resistance, membrane capacitance, resting membrane potential, and firing pattern) as described previously^1^.

### Pharmacology

**Table 1.**
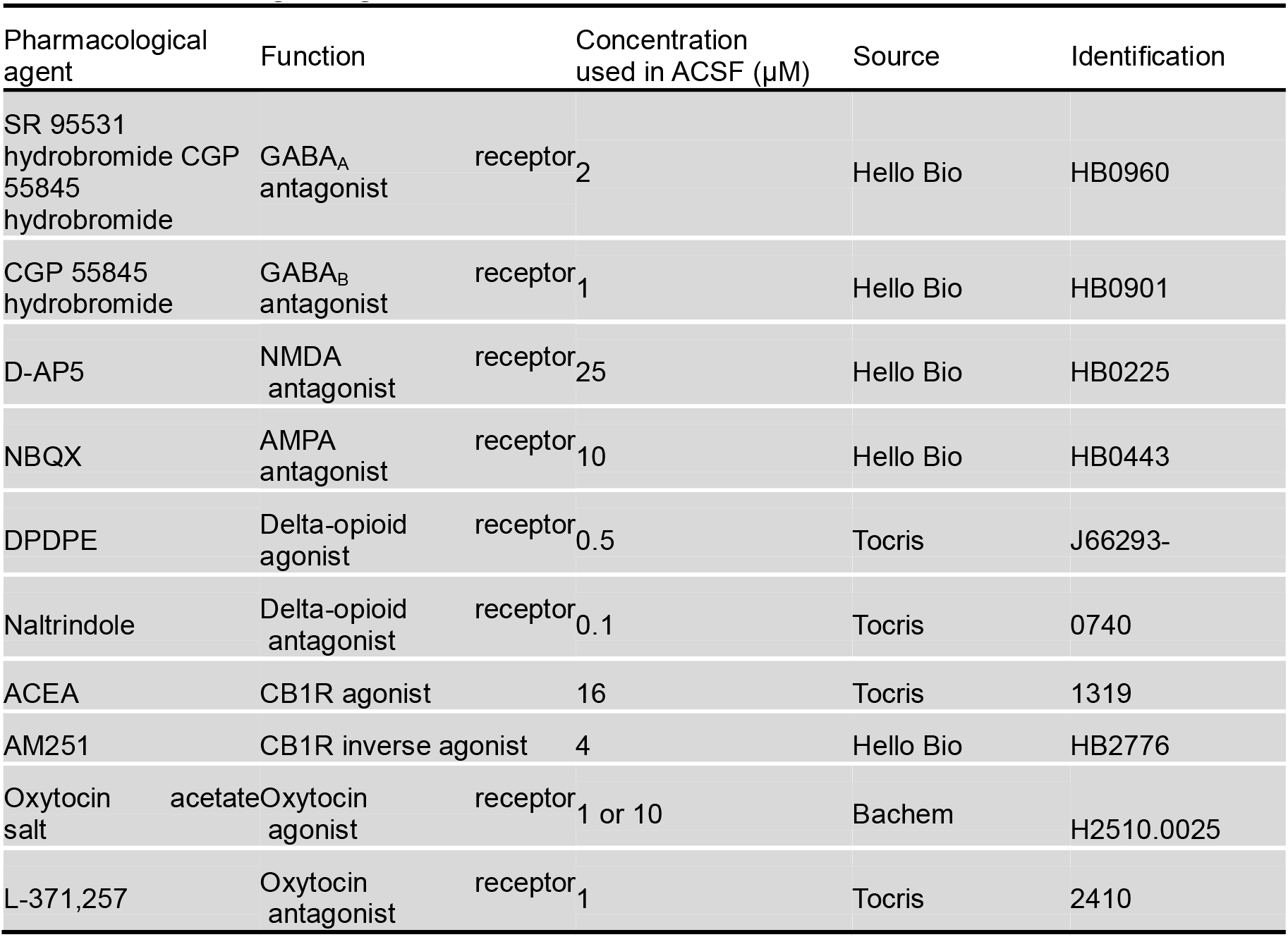
Pharmacological agents, sources and concentration.

#### Histology

After successful whole-cell recordings, slices were fixed in 4% paraformaldehyde (PFA) in phosphate buffered saline (PBS) at 4°C. Post-hoc labeling of filled cells with streptavidin-conjugated Alexa 488 was performed. Slices were permeabilized with 0.2 % Triton in PBS and blocked overnight with 3 % goat serum in PBS with 0.2 % triton. Slices were incubated overnight at 4°C in in 3 % goat serum in PBS, containing streptavidin-conjugated Alexa 488 (Life technologies, used at a 1:300 dilution) and Neurotrace Nissl (Life technologies, used at a 1:300 dilution). Slices were then washed in PBS and mounted. Slices were mounted with Pro-Long Diamond (Thermofisher) and z series confocal images were collected with a Zeiss 710 LSM microscope. Image analysis was performed with ImageJ and cellular morphologies were reconstructed from confocal images.

#### Biochemical measurements

Mice were divided into two groups: homecage or social interaction. Mice in the homecage group were left undisturbed until sacrificed. Mice in the social interaction group interacted with a novel mouse for 10 minutes and sacrificed 60 minutes after interaction. Finally, brains were extracted and fast-froze in liquid isopentane at -70°C. Samples were kept at -70°C until analyzed. Frozen brains were first mounted on the cryostat stage (Thermo Shandon, France) set at -15°C, then brains were sliced into 200μm-thick slices and bilateral CA1, CA2 and DG punches were collected on each slice and kept on carbon ice until storage at -70°C. RNA from brain punches was extracted using Trizol Reagent^TM^ (Thermofisher Scientific), then extracted RNA underwent several cleaning cycles with ethanol before storage at -20°C. Purity and concentration of total RNA were assessed by absorbance readings (Ratio A260/A230 and A260/A280) using the NanoDrop ND-1000 spectrophotometer (Thermo Fisher Scientific, Wilmington, DE). Then, RNA was reverse transcribed into cDNA using random primers and Transcriptase inverse SuperScript IVTM (Fisher scientific).

Gene amplification was performed using an SYBR Green I Master kit (ref RR420L, TAKARA BIO Europe, France) and appropriate custom primers. Primers for OTR, Wfs1, Rgs14, Actin, Gapdh and Rpl19 were designed and purchased from Eurofins Genomics (France; see below). Amplification reactions were run in duplicate using the LightCycler 480 system (Roche Diagnostics). First, primers accuracy was validated with a standard curve of four serial dilution points of whole hippocampus cDNA pool (ranging from 1000 pg to 1 pg of total RNA reverse transcripts), and a no template control (NTC). Tm calling curves were examined to verify the specificity of the amplified genes. To estimate CA1 and CA2 enrichment after punches, specific markers of the areas were quantified, using respectively Wfs1^2^ and Rgs14^3^. Three internal control genes, Actin, Gapdh and Rpl19 were quantified for accurate normalization of data. Normalization to reference genes and relative quantities determination were done using GenEx^TM^MultiD software.

OTR: AAAGGAAGGCTAGGGCTAACAGA(F), AGTGCCTATTTGCAGCCTTCA(R)

Wfs1: TCAGCAGTGAATCCAAGAACTACAT(F), GAATGATGCCCTTGGCGTACT(R)

Actin: AGTACGATGAGTCCGGCCCCT(F), AACGCAGCTCAGTAACAGTCCGC(R)

Gapdh: GAACATCATCCCTGCATCCA(F), CCAGTGAGCTTCCCGTTCA(R)

Rpl19: CCCGTCAGCAGATCAGGAA(F), GTCACAGGCTTGCGGATGA(R)

#### Reconstruction and analysis of PNs

PNs were reconstructed using Neurolucida software 360 (MBF Bioscience 2022) using the user-guided “directional kernels” algorithm. Neurolucida Explorer (MBF Bioscience 2022) was used in order to extract quantitative measures. Cell soma location was measured in order to make sure one group did not have a bias in the radial axis.

Sholl analysis (an analysis counting the number of dendritic intersections with a progressively larger concentric circle around the cell soma) was also accomplished through Neurolucida Explorer. 10µm steps were used for the sholl analysis of both basal and apical dendrites. Individual neuron sholl analyses were exported to Python in order to regularize by cell total length percentage (as cell length was highly dependent upon slice mounting) and average plots. This was done by taking the largest cell soma-dendritic end length as reference and doing a linear interpolation of missing data points for the smaller cells.

#### DREADD viral injections into hippocampal area CA2 and CNO injection

After 7–8 weeks under CD or HFD, *Cacng5*-Cre mice were anesthetized with air/isoflurane (4.5% induction; 1.5% maintenance) at 1L/min, injected with the analgesic buprenorphine (Buprécare, 0.1 mg/kg s.c. in 0.9% saline) and the non-steroidal anti-inflammatory drug carproxifen (Carprofène, 20mg/kg s.c. in 0.9% saline) and were placed on a stereotaxic apparatus (David Kopf Instruments). The scalp was shaved, disinfected with 10% povidone iodine and locally anaesthetized with a subcutaneous injection of lidocaine (Lurocaine 20mg/mL, 0.1ml non diluted). An anterograde adeno-associated virus carrying Cre-dependent inhibitory DREADD (AAV9/2-hSyn1-dlox-hM4DGi-mCherr) or control virus (AAV9/2-hSyn1-dlox-mCherry) was infused bilaterally using a 10 μL Hamilton syringe (Hamilton) and an ultra-micro pump (µMP3, World Precision Instruments, USA) over 2 min (100 nL/min) in the dorsal CA2 in each hemisphere (AP -1.5 mm, ML ± 2.4 mm, DV -1.8mm relative to Bregma). The pipette was left in place for a 5-minute diffusion period, before being slowly removed. The incision was closed with sutures, and the animal was kept on a heating pad until recovery. Mice were single housed for 2 days and their body weight and behavior were closely monitored during the 4 days following the procedure. Then, they were housed in groups of 4 to 8 mice per cage, and behavioral tests started 4 weeks later, enabling optimal virus expression.

To decrease the activity of CA2 PNs during social memory encoding, mice received an i.p. injection of the exogenous DREADD ligand Clozapine-N-oxyde (CNO, 2 mg/kg dissolved in 0.9% saline) 45 min before the sociability session. After behavior, mCherry expression in CA2 was verified histologically.

#### Intra-CA2 cannulas implantation and injection procedure

Stainless steel guide cannulas (28-gauge, Plastics One) were implanted bilaterally 1 mm above the CA2 (AP -1.5 mm; ML ±2.4 mm; DV -0.8 mm from Bregma). Guide cannulas were maintained in position with dental cement and dummy cannulas were always kept in each guide except during micro-infusions. Mice were allowed at least 5 days to recover from surgery, during which they were handled and weighed daily. The OT (Sigma–Aldrich) was dissolved in saline (0.9% NaCl) to obtain a final concentration of 1ng or 10ng /0.2 µL, corresponding to a concentration of 5µM and 50µM. The vehicle solution was saline. OT or saline was infused by inserting a 33-gauge internal cannula into the guide cannula. The internal cannula was connected to a 1µL glass syringe attached to an infusion pump (Harvard Apparatus) and projected an additional 1 mm ventral to the tip of the guide cannula. A total volume of 0.2µL was delivered bilaterally at a rate of 0.1 µL/min. The internal cannula remained in place for a further 1 min after the infusions and was then removed. The concentration, rate and infusion volumes were chosen based on previous literature^4–6^.

All mice were sacrificed after behavioral testing. Animals were deeply anaesthetized with a mix of pentobarbital and lidocaine (Exagon, 300 mg/kg and Lurocaine, 30 mg/kg) before cardiac perfusion with phosphate-buffered saline solution followed by 4% paraformaldehyde (Sigma-Aldrich) diluted in phosphate-buffered saline. Brains were collected and post-fixed in 4% paraformaldehyde at 4°C, then stored at 4°C until slicing. 40μm coronal sections were cut on a vibratome (Leica) and stored in cryoprotective solution (glycerol and ethylene glycol; Sigma-Aldrich) at −20°C. Every sixth section was mounted on gelatin-coated slides, covered by medium (Southern Biotech) and cover-slipped. The slices were visualized under a microscope and each section was photographed (Nikon-ACT-1 software), enabling assessment of correct cannula implantation.

#### Test for sociability and SRM

CD and HFD-fed mice behavior was assessed in an open-field Plexiglas arena (45 × 25 cm and 20 cm high) placed in a sound attenuated room adapted from ^7,8^. The assay paradigm comprises three sessions: habituation, training, and test. In the first session, mice were placed in the arena containing two empty pencil-wire cups positioned on opposing sides and left free to explore for 10 min. Then, 24 h after this session, a juvenile mouse (stimulus, 6−7 weeks old), which had no prior contact with the subject mice, was placed under one of the wire cups while the other cup remained empty. The subject mouse was then placed in the arena and was left free to explore it for 10 min. During the third session, performed either 15 minutes or 3 hours later, the same stimulus animal was again placed under the wire cup and a novel unfamiliar juvenile mouse was placed under the opposing cup. Experimental mice were then placed again in the arena and tested for discrimination between novel and familiar mouse in a 10 min session. Each experimental mouse was subjected to the procedure separately, and care was taken to remove any olfactory/taste cues by cleaning carefully the arena and wire cups between trials. The positions of the social stimuli (empty × social; familiar × novel) were counterbalanced across subjects and trials to prevent bias due to place preference. All juvenile stimulus mice were habituated to remain under the wire cups for 30 min during several days before behavioral testing. The animal’s behavior during all sessions was videotaped, and the time spent actively exploring the stimuli was analyzed by experienced observers unaware of the experimental groups. Exploration was defined as direct snout-to-cup contact, and the time spent climbing on the cups was not considered. Data are expressed as a percentage of time spent exploring each cup (social × non-social during the second session or familiar × novel during the third session). The raw exploration time data were also employed to determine social and discrimination indexes, according to the following equations: Sociability index = (time exploring social cup-time exploring non-social cup)/ total exploration time of both cups; Discrimination Index = (time exploring novel mouse-time exploring familiar mouse)/ total time of exploration of both mice).

#### Test for social discrimination between cage-mate and novel mouse

The assay paradigm was similar to the one used for SRM but comprises only two sessions: habituation and discrimination test. In the first session, mice were placed the arena containing two empty pencil-wire cups positioned on opposite sides and left free to explore for 10 min. Then, 24 h after this session, the subject mouse was placed in the arena again and left free to explore for 10 min two mice placed under the wire cups. One mouse (novel) never had contact with the subject and the other mouse was a familiar co-housed littermate of the subject mouse (cage-mate). Novel and cage-mate stimulus mice were habituated to remain under the wire cups for 30 min the day before behavioral testing. Data are expressed as Discrimination Index = (time exploring novel mouse-time exploring cage-mate)/ total time of exploration of both mice).

#### Object recognition memory

During training, two identical new objects were presented in a novel open field arena (40 × 40 × 40 cm, wood) and each mouse was left freely to explore them. Long-term memory was assessed 24 hours later, by randomly replacing one of the objects by a novel one. In both phases, object exploration, defined as nose and whiskers pointed towards the object in a distance of less than 1–1.5 cm away, was quantified by a trained experimenter blind to experimental groups. In both phases, the session duration was 10 min max but the mouse was removed from the apparatus after 20 s of total exploration for both objects which reduces inter-individual variability ^9^, otherwise mice were excluded from analysis. Data are presented as the percentage of exploration of novel object over total exploration time of both objects.

## QUANTIFICATION AND STATISTICAL ANALYSIS

### AP shape analysis

AP shape analysis was always performed on the first AP elicited at rheobase. Axograph was used to measured AP amplitude, 20-80% rise-time, half-width (the width of the AP at half of its maximum amplitude), decay time-constant (measured as the time of 67% AP decay) and after-hyperpolarization (measured as the minimal potential after each AP subtracted from the cell potential before AP firing) for each spike.

### Burst analysis

Bursts were defined as groups of at least 2 action potentials firing at a frequency of more than 0.5Hz, lasting at least 1.5 seconds, and characterized by a slow rampant depolarization as well as an after-hyperpolarization (as is observed on the example shown in figure 3E). Bursting onset (measured as the time from drug application to first burst), Burst duration (measured as the duration of the depolarizing plateau phase of the burst), firing rate during bursts, number of APs per burst, inter-burst intervals and total number of bursts elicited (calculated withing 15 minutes of drug application) were measured via Axograph.

### Statistics

Origin Pro software was used for electrophysiology data analysis and statistics. All statistical details can be found in the figure legends and results. For all electrophysiology experiments, the number of slices and number of animals are indicated. For behavioral experiments, the number of animals is indicated.

The Shapiro-Wilk test was used to determine whether the data followed a normal distribution and an F-test to see if was homoscedastic. We used Student’s t tests (one-sample, paired or unpaired) and 2-way ANOVAs when the data followed a normal distribution and were of equal variance. When the distribution was not normal or the variances unequal, we used non-parametric tests (Mann-Whitney, Kruskal-Wallis U-test). Statistical significance was set to p < 0.05 (*** indicates p < 0.001, ** indicates p < 0.01, and * indicates p < 0.05). All values are reported as the mean ± SEM.

