## Supplementary material for "Obesogenic Diet Impairs Social Memory Through Alterations of Hippocampal CA2 Excitability and Oxytocin Signaling": Total Supplemental

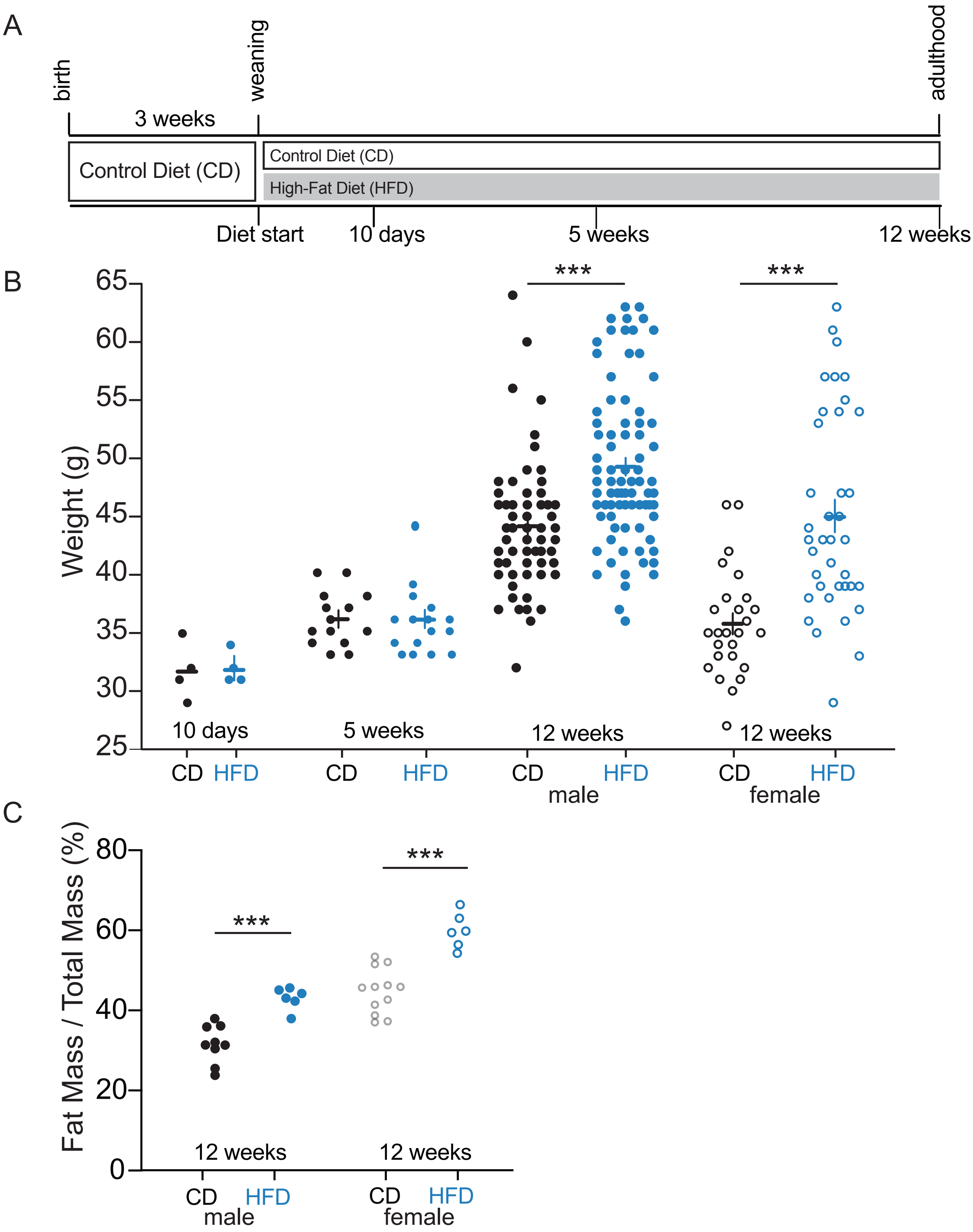


**Supplemental Figure 1.** A long-term, but not short-term, high-fat diet induces weight gain in male and female mice. A. Experimental timeline. B. 10 days or 5 weeks of a HFD did not significantly increase weight of animals (HFD: n=4 and CD: n=4 for at 10 days, statistics NA; HFD: n=16, CD: n=14 for 5 weeks of diet; two-sample t-test p=0.74). Right, 12 weeks of a HFD increased the weight of male and female mice (CD: n=58, HFD: n=80, two-sample t-test p=5.2E-6 for males; CD: n=26, HFD: n=38, two-sample t-test p=4.5E-6 for females). C. A 12-week HFD increases the fat-mass of male and female mice (CD: n=9; HFD: n=6 for males, two-sample t-test p<0.001; CD: n=12. HFD: n=6 for females, p<0.001). Filled circles represent male mice; empty circles represent female mice.

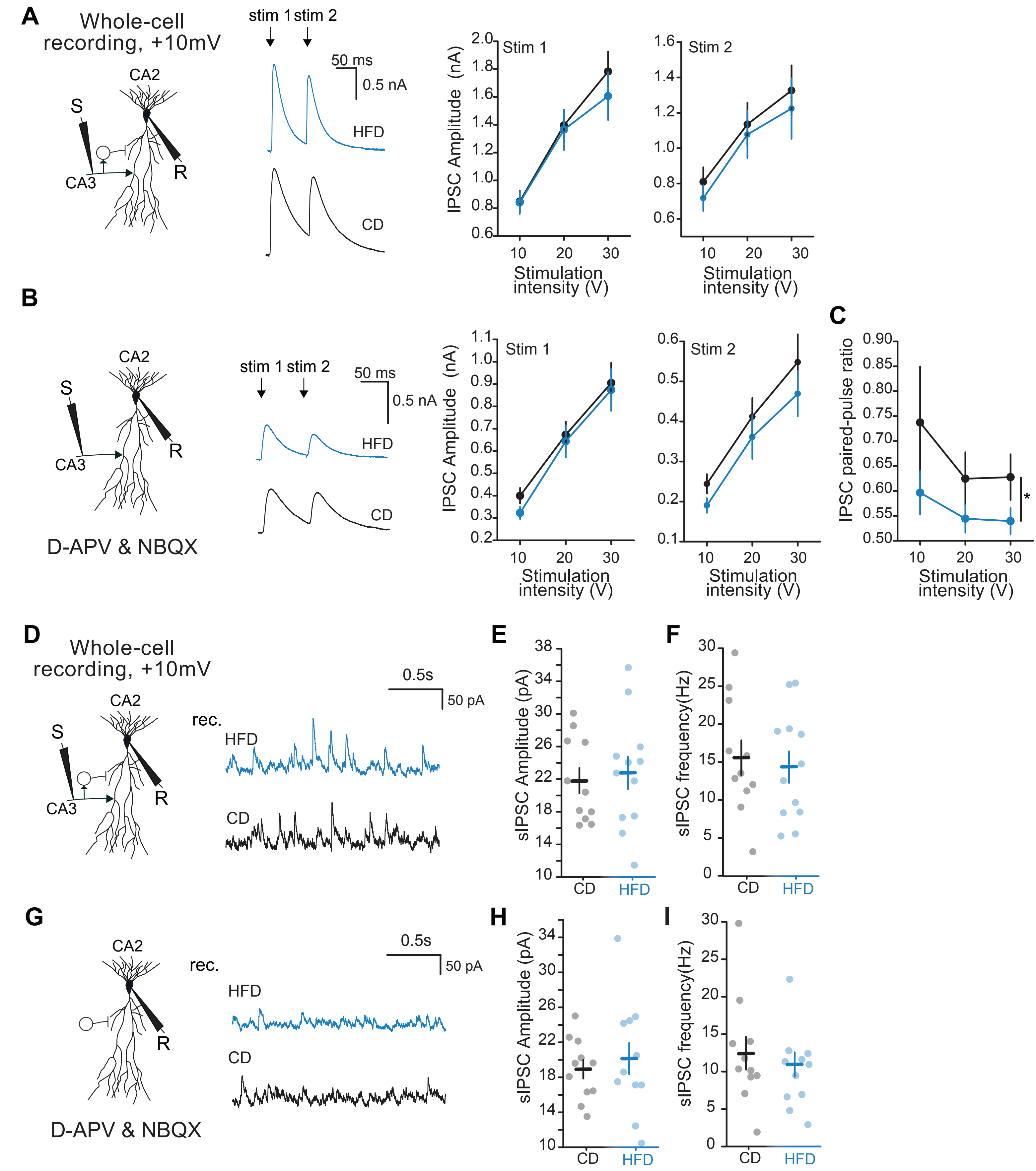


**Supplemental Figure 2.** Effect of HFD-exposure on inhibitory transmission. A. Left: schematic of recording conditions. Middle: example traces of total inhibitory transmission onto CA2 PNs upon SC stimulation. Right: A HFD does not alter total evoked IPSC amplitude onto CA2 PNs upon SC stimulation (CD: n=18,11; HFD: n=17,12; Two-way ANOVA p=0.086 for diet effect of stimulation 1; p=0.41 for diet effect of stimulation 2). B. Left: schematic of recording conditions. Middle: example traces of direct inhibitory transmission onto CA2 PNs upon SC stimulation. Right: HFD exposure does not alter evoked direct IPSC amplitude onto CA2 PNs upon SC stimulation (CD: n=18,11; HFD: n=17,12; Two-way ANOVA p=0.40 for diet effect of stimulation 1; p= 0.12 for diet effect of stimulation 2). C. HFD exposure decreases the inhibitory paired pulse ratio onto CA2 PNs (CD: n=18,11; HFD: n=17,12; Two-way ANOVA p= 0.041 for diet effect). D. Left: schematic of recording conditions. Right: example traces of total spontaneous inhibitory transmission onto CA2 PNs. E. HFD exposure does not affect total spontaneous IPSC amplitude (CD: n=12,7; HFD: n=12,7; two-sample t-test p= 0.71). F. HFD exposure does not affect total spontaneous IPSC frequency (CD: n=12,7; HFD: n=12,7; two-sample t-test p= 0.69). G. Left: schematic of recording conditions. Right: example traces of direct spontaneous inhibitory transmission onto CA2 PNs. H. HFD exposure does not affect direct spontaneous IPSC amplitude (CD: n=12,7; HFD: n=12,7; two-sample t-test p= 0.57). I. A HFD does not affect direct spontaneous IPSC frequency (CD: n=12,7; HFD: n=12,7; two-sample t-test p= 0.58).

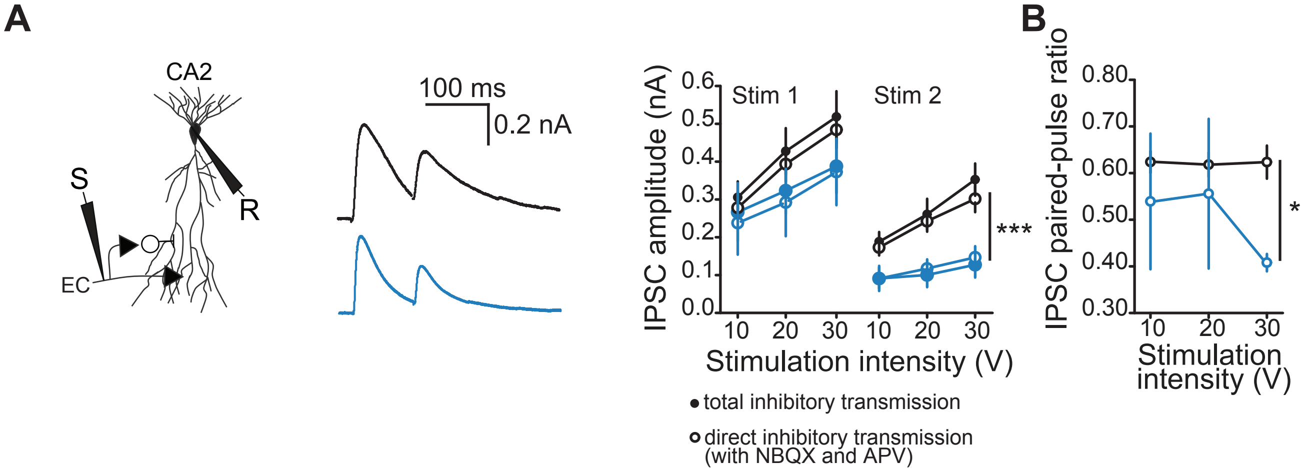


**Supplemental Figure 3.** A HFD decreases inhibitory transmission at EC-CA2 synapses. A. left, Schematic of recording conditions. Middle: example traces of inhibitory transmission onto CA2 PNs upon PP stimulation. Right: A HFD decreases IPSC amplitude in CA2 PNs upon PP stimulation (CD: n=8,6; HFD: n=5,4; Two-way ANOVA p= 0.087 for diet effect of stimulation 1 without blockers of excitatory transmission; p= 0.093 for diet effect of stimulation 1 with blockers of excitatory transmission; Two-way ANOVA p = 1.2E-5 for diet effect of stimulation 2 without blockers of excitatory transmission; p =1.6E-5 for diet effect of stimulation 2 with blockers of excitatory transmission). B. A HFD decreases the direct inhibitory EC-CA2 paired pulse-ratio (CD: n=8,6; HFD: n=5,4; Two-way ANOVA p= 0.048 for diet effect).

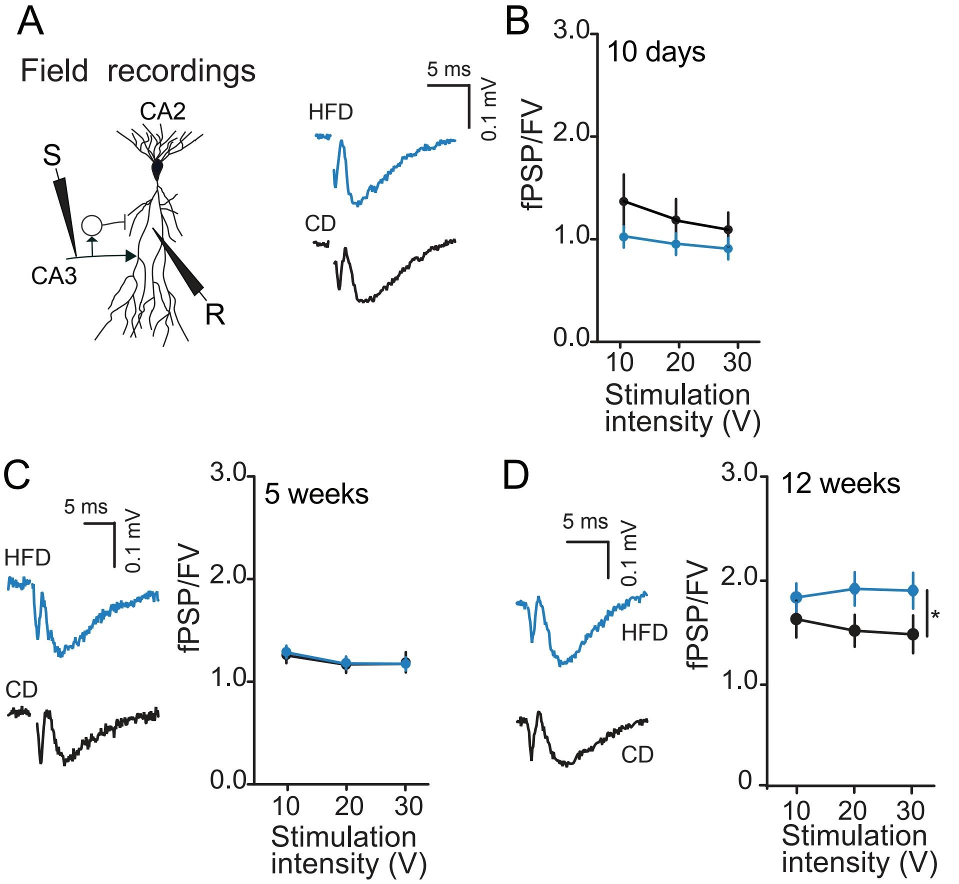


**Supplemental Figure 4.** A long-term but not short-term exposure to HFD increases CA3 to CA2 transmission. A. Schematic of recording condition. B. A 10-day HFD exposure does not alter CA2 fPSPs normalized to the FV upon SC stimulation (CD: n=12,4; HFD: n=12,4, Two-way ANOVA p=0.071). C. A 5-week HFD exposure does not alter CA2 fPSPs normalized to the FV upon SC stimulation (CD: n=23,6; HFD: n=31,8, Two-way ANOVA p=0.83). D. A 12-week HFD increases CA2 fPSPs normalized to the FV upon SC stimulation (CD: n=13,6; HFD: n=23,10, Two-way ANOVA p=0.016 for diet effect).

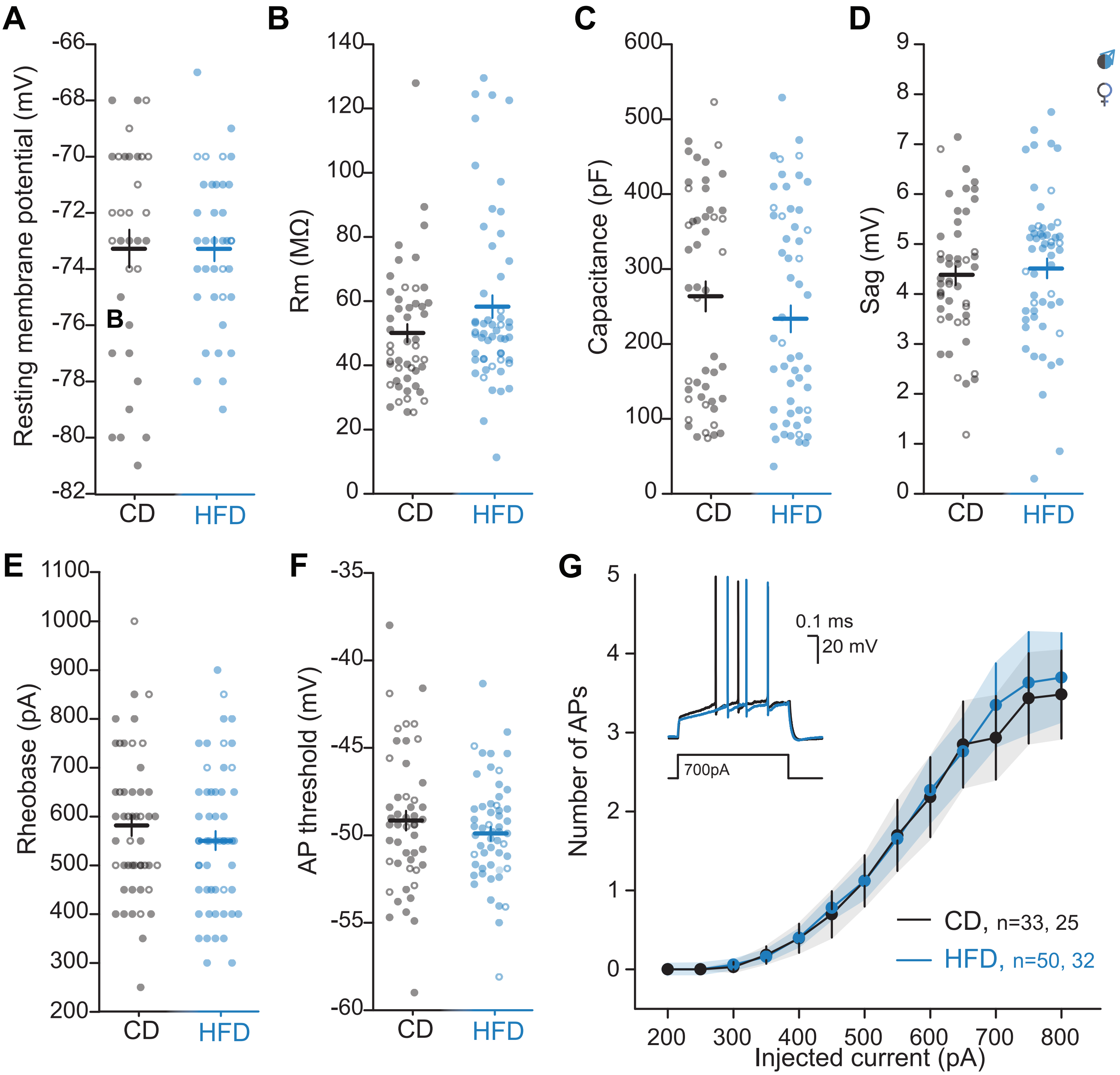


**Supplemental Figure 5.** Effect of a HFD on intrinsic properties and excitability of CA2 PNs. A. Resting membrane potential (CD: n=33,24; HFD: n=37,26; Mann-Whitney U-test p= 0.56). B. Membrane resistance (CD: n= 49,36; HFD: n=54,35; two-sample t-test p= 0.052.). C. Membrane capacitance (CD: n= 49,36; HFD: n=54,35; two-sample t-test p= 0.35). D. Sag potential at -100 mV during a 1-second hyperpolarizing current injection (CD: n= 49,36; HFD: n=54,35; two-sample t-test p= 0.51). E. Rheobase (CD: n= 49,36; HFD: n=54,35; two-sample t-test p= 0.33). F. Action potential threshold (CD: n= 49,36; HFD: n=54,35; two-sample t-test p= 0.48). G. The number of APs elicited as a function of current injection (CD: n= 49,36; HFD: n=54,35; Two-way ANOVA p=0.78 for diet effect). Filled circles represent data from male mice; empty circles data from female mice.

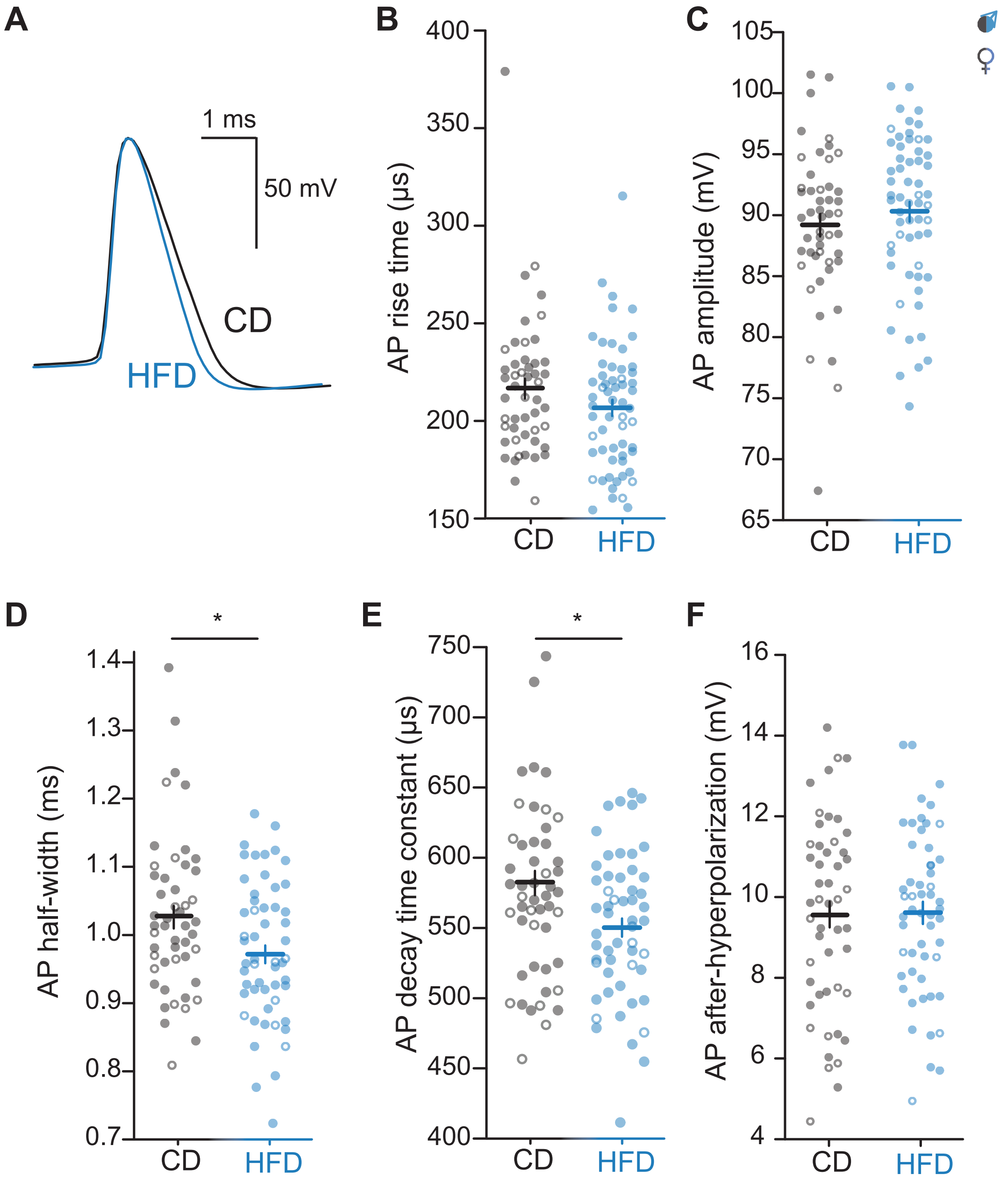


**Supplemental Figure 6.** A HFD shortens CA2 PN action potentials. A. Example shape of an action potential of a CA2 PN for CD and HFD-fed mice. B. AP rise time (CD: n=49,36; HFD: n=54,35; two-sample t-test p= 0.22). C. AP amplitude (CD: n=49,36; HFD: n=54,35; two-sample t-test p= 0.47). D. AP halfwidth was significantly reduced in HFD conditions (CD: n=49,36; HFD: n=54,35; two-sample t-test p= 0.020). E. AP decay time-constant was significantly decreased in HFD conditions (CD: n=49,36; HFD: n=54,35; two-sample t-test p= 0.035). F. AP after-hyperpolarization (CD: n=49,36; HFD: n=54,35; two-sample t-test p= 0.88). Filled circles represent CA2 PNs from male mice; empty circles represent CA2 PNs from female mice.

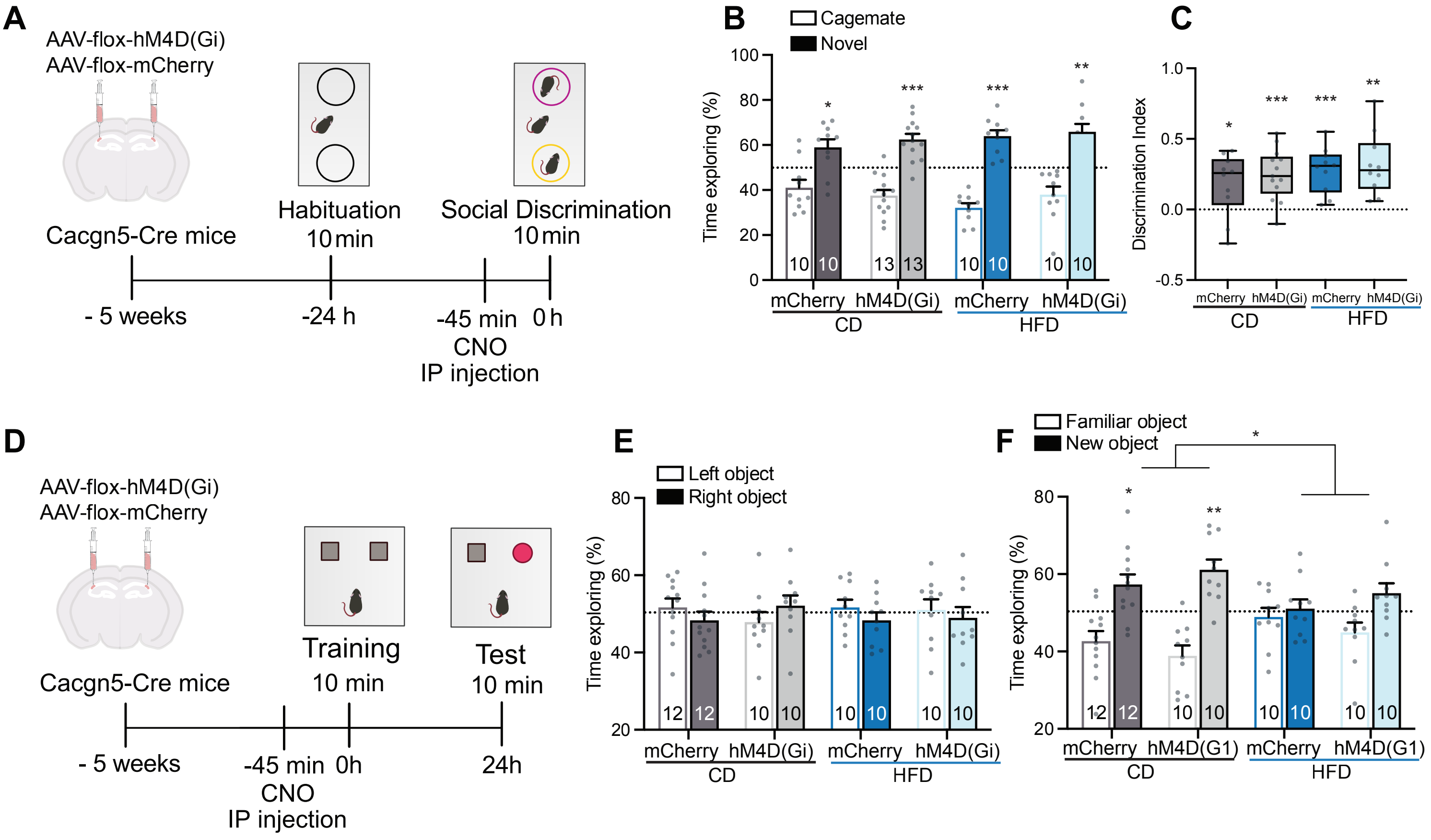


**Supplemental Figure 7** A. Behavioral paradigm of AAV-mediated Gi-DREADD expression in CA2 PNs for SRM testing between a cagemate and novel conspecific. B. HFD and CD mice, injected with either mCherry control virus or Gi-DREADD virus, explore the novel mouse a greater amount of time compared to the cagemate ($p<0.05$ for CD-mCh, $p<0.005$ for CD-Gi and HFD-mCh, and $p<0.01$ for HFD-Gi comparison between the percentage of time exploring novel mouse and 50% using one-sample t-test).C. Discrimination index is higher than 0 for all conditions. D. Behavioral paradigm of AAV-mediated Gi-DREADD expression in CA2 PNs for novel object discrimination testing. E. No significant difference in exploration time of identical objects in different locations was observed for all conditions. F. In the test phase, the CD-nCh ($p < 0.05$) and CD-Gi ($p < 0.01$) mice showed increase exploration of the novel object, whereas the HFD-fed animals did not show significant exploration times of the novel and familiar object. The number of animals used per group is shown on the figure.

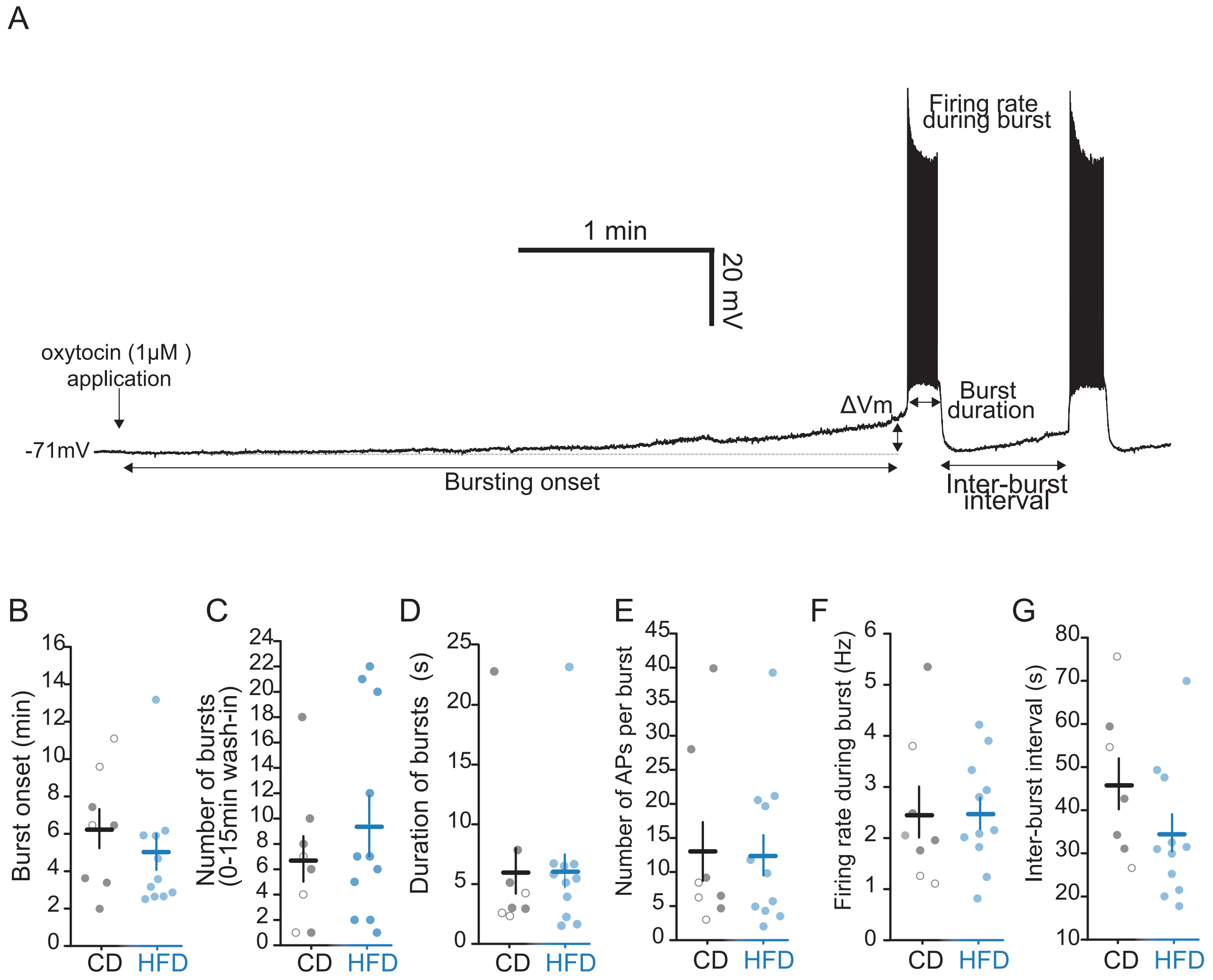


**Supplemental Figure 8.** Further Effects of HFD-exposure on OT-induced CA2 PN depolarization and bursting. A. Example trace of oxytocin-induced depolarization and bursting of CA2 PN with the different measurement indicated for comparison of bursting dynamics. B. OT-induced burst onset (CD: n=8,6; HFD: n=11,6; Mann-Whitney U-test p=0.17). C. OT-induced burst number (CD: n=8,6; HFD: n=11,6; Mann-Whitney U test p=0.56). D. OT-induced burst duration (CD: n=8,6; HFD: n=11,6; Mann-Whitney U test p=0.83). E. Number of APs per burst (CD: n=8,6; HFD: n=11,6; Mann-Whitney U test p= 0.96). F. Firing rate during burst (CD: n=8,6; HFD: n=11,6; two-sample t-test p=0.98). G. Inter-burst interval (CD: n=8,6; HFD: n=11,6; two-sample t-test p=0.1465). Filled circles represent CA2 PNs from male mice; empty circles represent CA2 PNs from female mice.

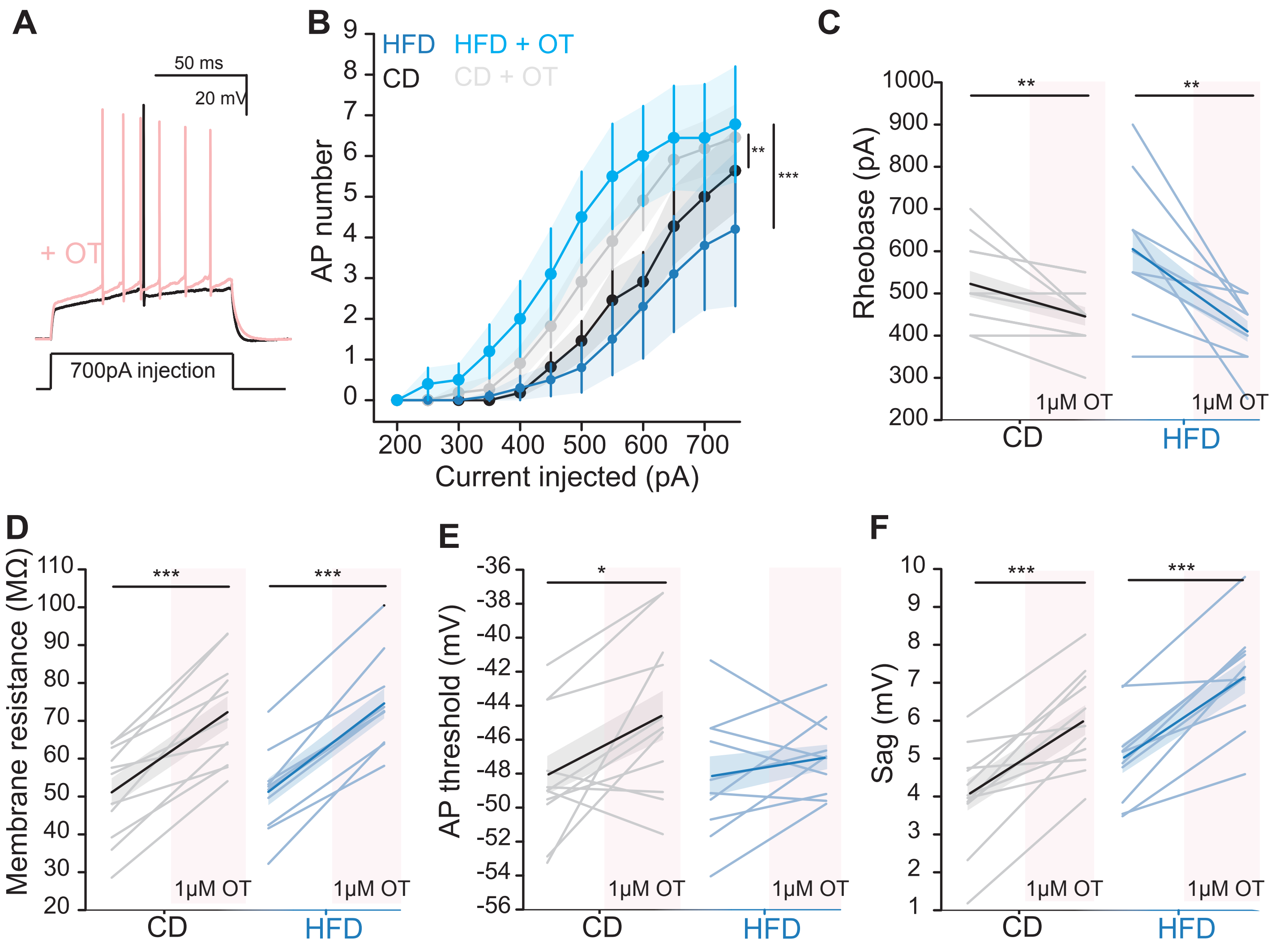


**Supplemental Figure 9.** A HFD does not modify OT-induced enhanced CA2 PN excitability. A. Example of CA2 PN depolarization and AP firing in response to positive current injection before and after 1µM OT application. B. AP number elicited by current injection (CD: n=11,7 Two-way ANOVA p=0.0078 before and after OT; HFD: n=10,7; p=2.5E-4 before and after OT). C. Rheobase current (CD: n=11,7; HFD: n=10,7; paired-sample t-test p=0.0085 for comparison before and after OT for CD; paired-sample t-test p=0.0028 for comparison before and after OT for HFD). D. Membrane resistance (CD: n=11,7; HFD: n=10,7; paired-sample t-test p= 2.5E-5 for comparison before and after OT for CD; paired-sample t-test p= 1.3E-6 for comparison before and after OT for HFD). E. AP threshold (CD: n=11,7; HFD: n=10,7; paired-sample t-test p= 0.021 for comparison before and after OT for CD; paired-sample t-test p=0.27 for comparison before and after OT for HFD). F. CA2 PN sag (CD: n=11,7; HFD: n=10,7; paired-sample t-test p=1.8E-4 for comparison before and after OT for CD; paired-sample t-test p=1.0E-4 for comparison before and after OT for HFD).

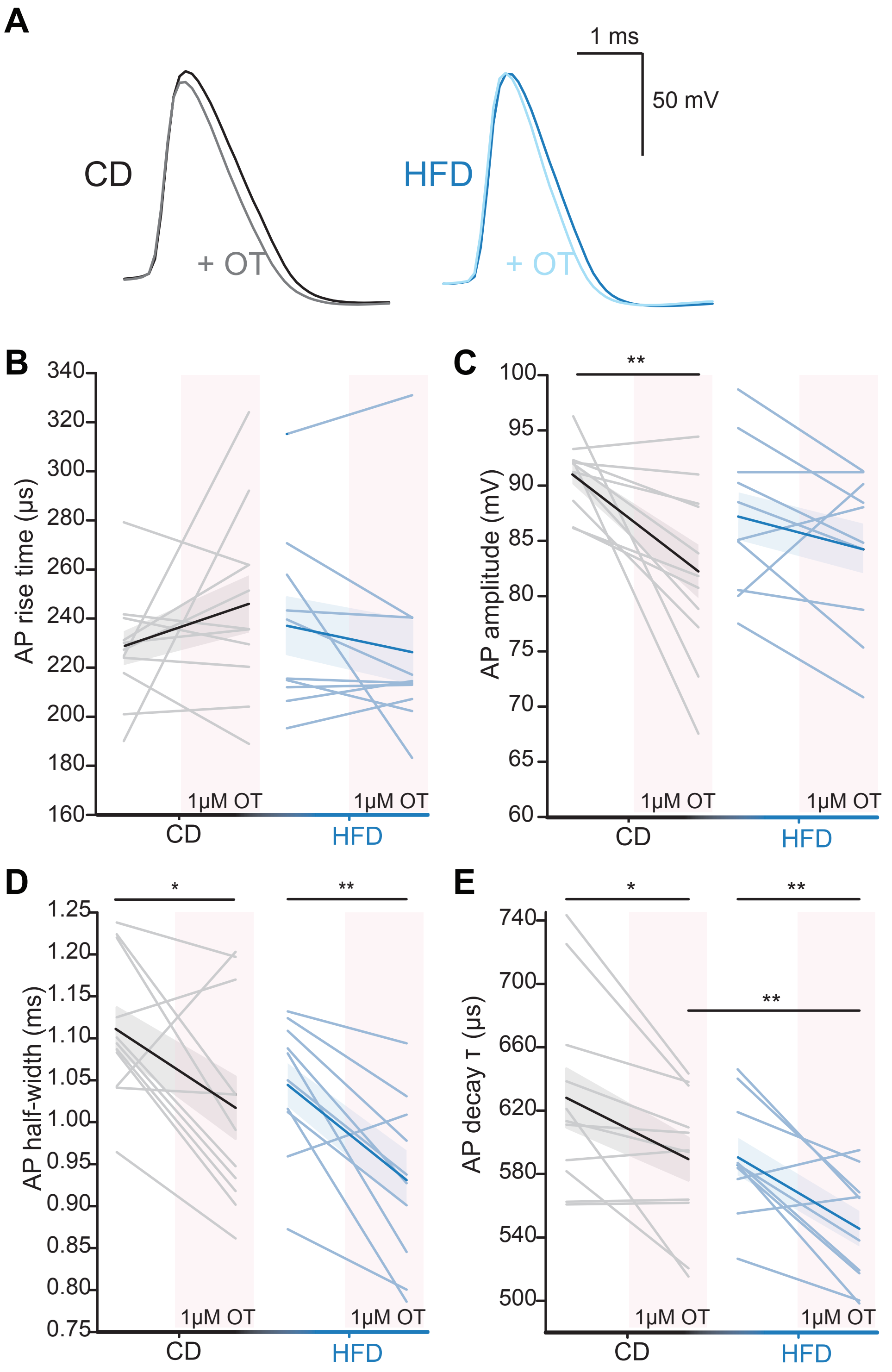


**Supplemental Figure 10.** 1µM OT application shortens CA2 PN action potentials in CD and HFD-fed mice. A. Example traces of CD and HFD CA2 PN action potentials induced by current injection, in the presence and absence of 1µM OT. B. AP rise-time in (CD: n=11,7; paired-sample t-test p=0.20 before and during OT; HFD: n=10,7; paired-sample t-test p=0.23 before and during OT). C. AP amplitude (CD: n=11,7; paired-sample t-test p= 0.0066 before and during OT; HFD: n=10,7; paired-sample t-test p= 0.15 before and during OT). D. AP half-width (CD: n=11,7; paired-sample t-test p= 0.026 before and during OT; HFD: n=10,7; p= 0.0023 before and during OT; p= 0.080 between CD OT and HFD OT). E. AP decay time-constant (CD: n=11,7; paired-sample t-test p= 0.013 before and during OT; HFD: n=10,7; paired-sample t-test p= 0.0039 before and during OT; paired-sample t-test p= 0.0064 between CD OT and HFD OT).

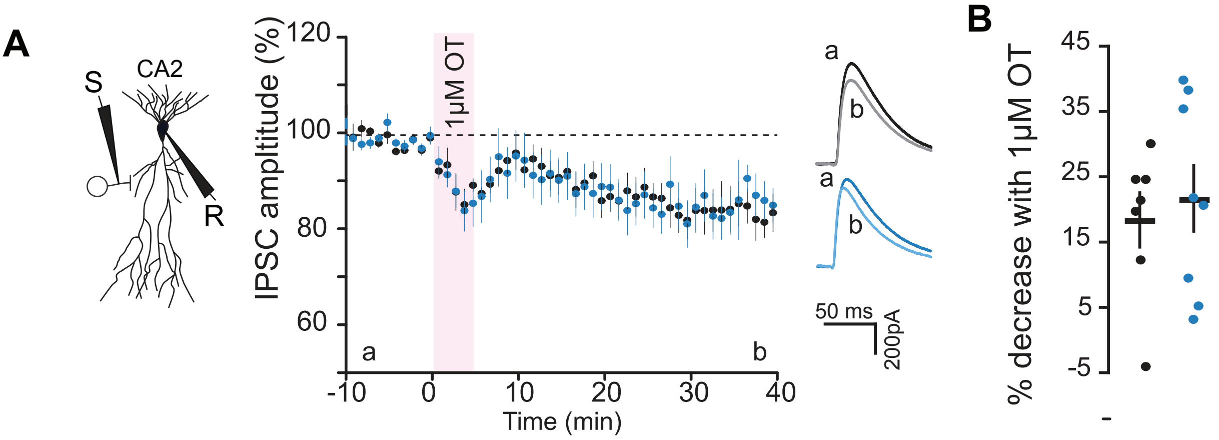


**Supplemental figure 11**. HFD exposure does not alter changes in inhibitory transmissioni following oxytocin application has no effect on OT-induced changes in inhibitory synaptic transmission (A) Left: Schematic of whole-cell recording of CA2 PN upon stimulation in SR, in the presence of excitatory transmission blockers. CA2 PNs were held at +10mV in voltage-clamp. Middle: 5-minutes 1µM OT application induces a long-term decrease in IPSC amplitude in CD and HFD-fed mice CA2 PNs. (B) Percentage of 1µM OT-induced IPSC depression for each CA2 PN (measured at 10 last minutes of recording normalized to baseline) (CD: n=7,5 one-sample t-test p=0.0049 baseline vs end; HFD: n=8,6, one-sample t-test p=0.0041 baseline vs end; CD vs HFD: unpaired t-test p= 0.63). Filled circles represent CA2 PNs from male mice; empty circles represent CA2 PNs from female mice.

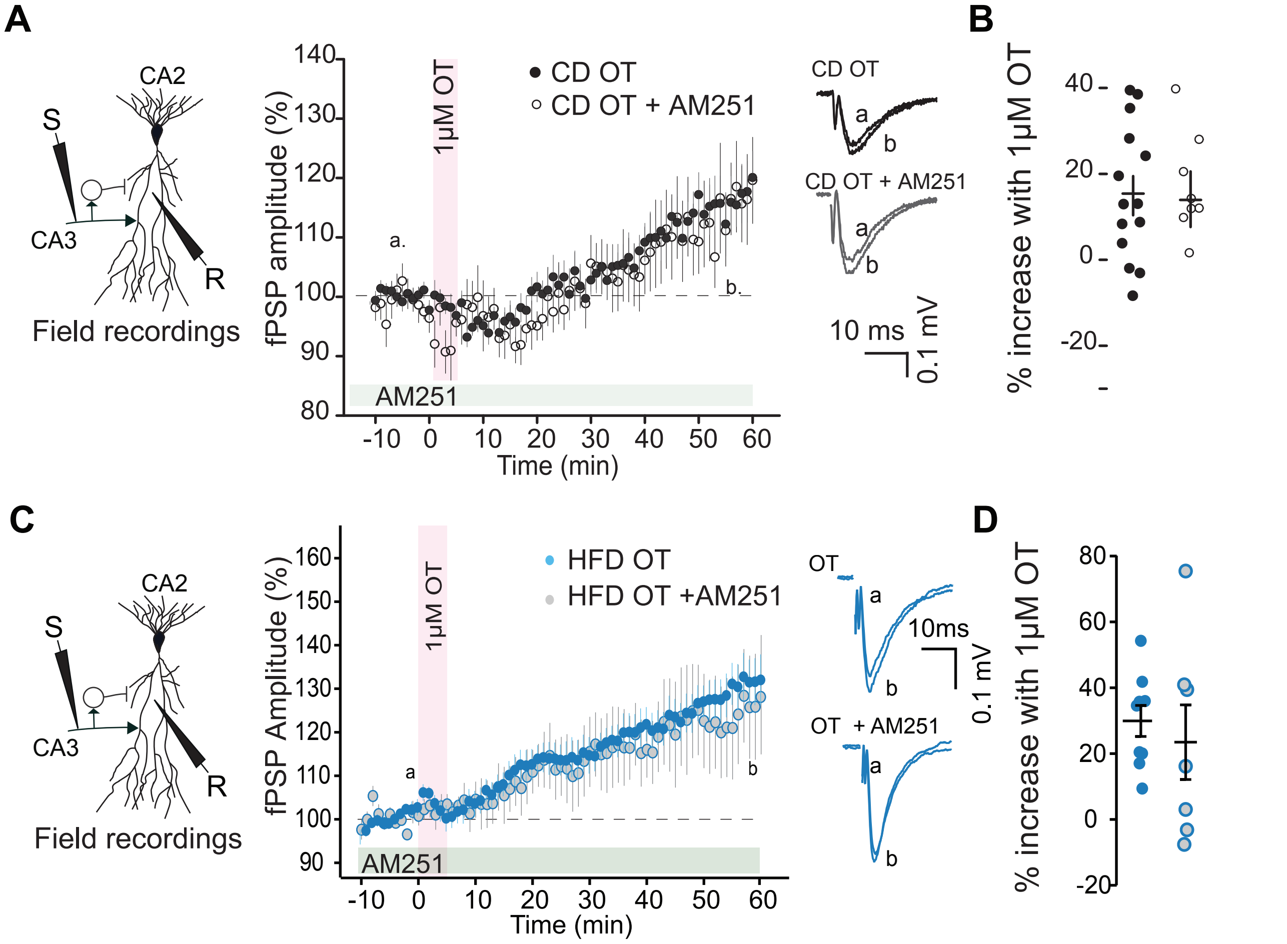


**Supplemental figure 12**. Oxytocin application has no effect on eCB-mediated synaptic plasticity in area CA2 in both CD and HFD conditions. (A) Left: Schematic of CA2 fPSP recording upon SC stimulation. Middle: 5-minuteminute 1µM OT application induces long-term increase of CA2 fPSP amplitude in CD--fed animals, with or without the presence of AM251 (CB1R antagonist). Right: example traces before and after OT application with and without AM251 application. (B) Percentage of 1µM OT-induced potentiation with and without the presence of AM251 (CB1R antagonist) for each slice recording (measured at 10 last minutes of recording normalized to baseline) (CD OT only: n=14,12, one sample t-test p=0.0021; CD OT+AM251: n=8,8, one-sample t-test p=0.062 for difference with baseline; CD OT only vs CD OT+AM251 two-sample t-test p=0.7) (C) Left: Schematic of CA2 fPSP recording upon SC stimulation. Middle: 5-minute 1µM OT application induces long-term increase of CA2 fPSP amplitude in HFD-fed animals, with or without the presence of AM251 (CB1R antagonist). Right: example traces before and after OT application with and without AM251 application. (D) Percentage of increase in the field potential following 1µM-OT application is similar with and without blocking CB1Rs (HFD OT only: n=9,33, one-sample t-test p= 0.0002; HFD OT+AM251: n=7,3, one-sample t-test p=0.084 for difference with baseline; HFD OT only vs HFD OT+AM251, two-sample t-test p=0.613) Filled circles represent CA2 PNs from male mice; empty circles represent CA2 PNs from female mice.

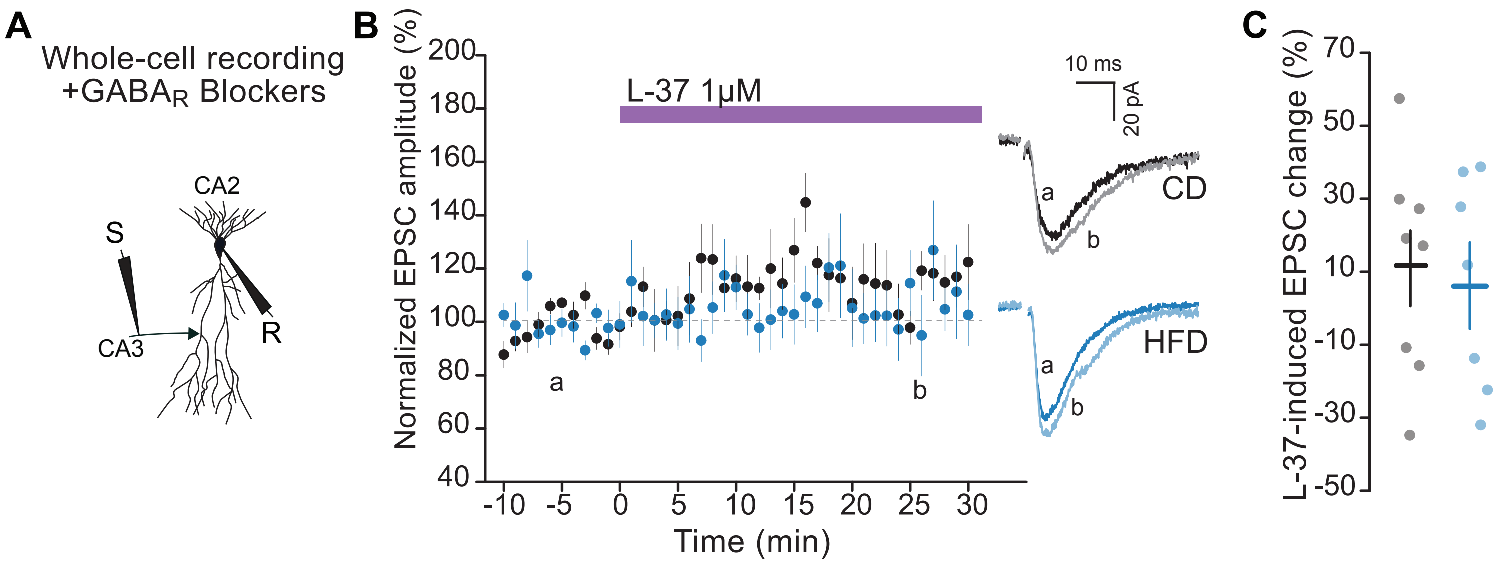


**Supplemental Figure 13.** Blocking OT receptors does not modify CA2-CA2 excitatory transmission. Left: Schematic of whole-cell EPSC recording in the presence of GABAA and GABAB receptor blockers. CA2 PNs were held at -70mV in voltage clamp mode. Middle: 1µM L-37 application did not modify CA3-CA2 excitatory transmission amplitude in either CD or HFD-fed mice. Example traces of CA2 PN EPSPs from CD and HFD-fed mice. Right: L-37-induced EPSC change for all CA2 PNs recorded (CD: n=8,6 one-sample t-test p=0.27 baseline vs end; HFD: n=7,4 one-sample t-test p=0.65 baseline vs end; CD vs HFD: two-sample t-test p= 0.88).

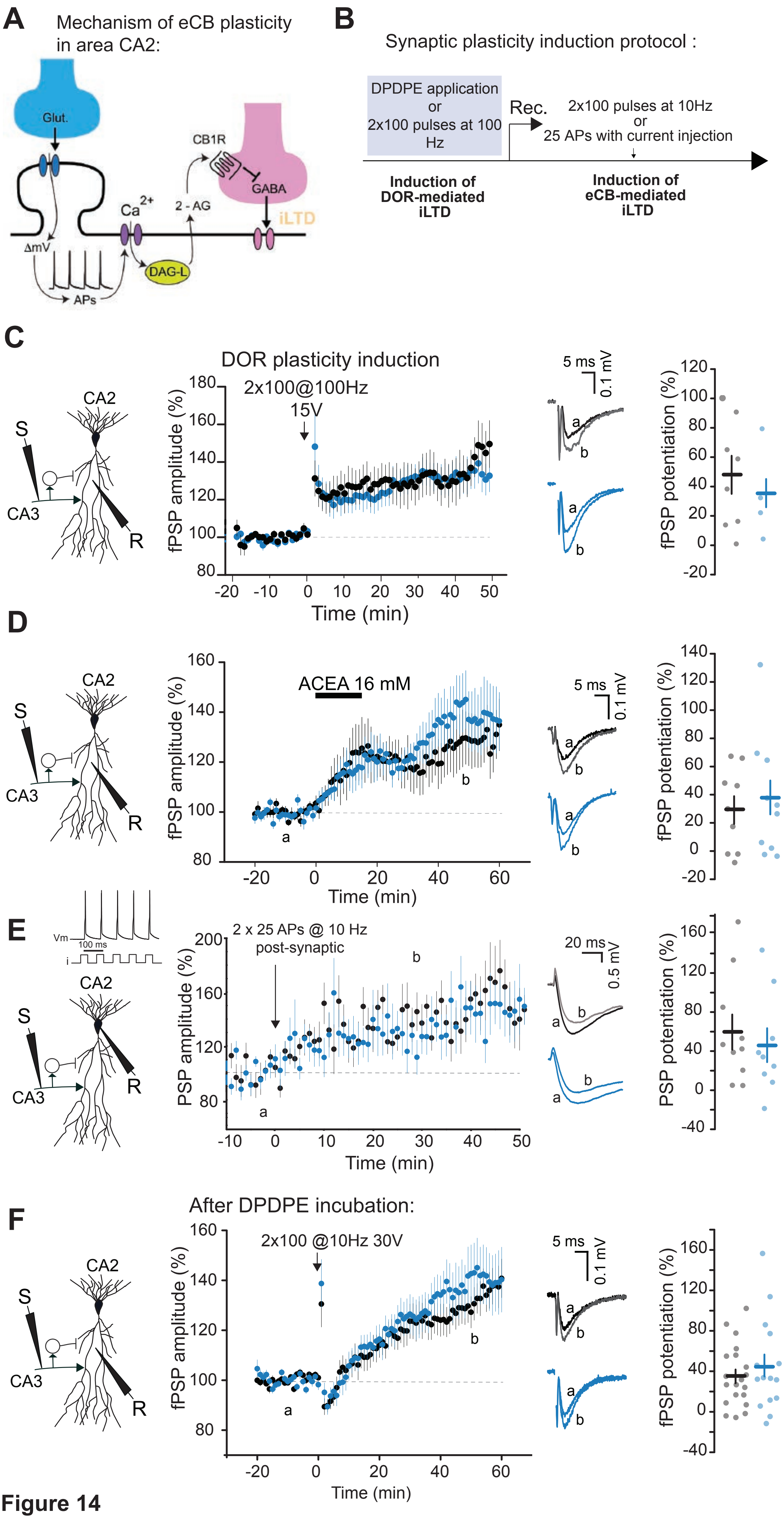


Supplemental Figure 14

**Supplemental Figure 14.** A HFD does not alter eCB-mediated plasticity of inhibitory transmission in area CA2. A. Cartoon summarizing the molecular mechanism of eCB plasticity at inhibitory synapses in area CA2. Post-synaptic depolarization, presumably resulting from excitatory synaptic transmission, leads to AP firing. This AP firing leads to opening of voltage-activated calcium channels and calcium influx. Calcium influx leads to the synthesis of 2arachidonoylglycerol (2-AG) by diacylglycerol lipase (DAG-L). 2-AG is the endogenous ligand for cannabinoid type 1 receptors (CB1R). Activation of pre-synaptic CB1Rs results in long-term depression of GABAergic transmission. B. Experimental protocol portraying how the DOR-mediated and ECB-mediated plasticities are sequentially induced in area CA2. Slices were pre-incubated in DPDPE, a ligand or delta opioid receptors (DOR), or subjected to a high-frequency (100Hz) stimulation protocol. Activation of DORs results in a long-term depression of inhibitory transmission at parvalbumin-expressing inhibitory terminals, and allows CA2 PNs to be effectively depolarized by glutamatergic transmission (Loisy et al., 2022). C. Left: Schematic of extra-cellular recording configuration. Middle: The DOR-mediated plasticity in area CA2 is unaltered by HFD exposure. A high frequency induction protocol of 100 stimulations at 100 Hz, repeated twice induces a long-term increase of fPSP amplitudes in both CD and HFD-fed mice. Right: 100 Hz-induced fPSP increase is similar in HFD and CD-fed mice (CD: n=8,4; HFD: n=11,4; two-sample t-test p= 0.44). D. Left: Schematic of extracellular recording configuration. Middle: 15 minutes of 16 mM ACEA application induces a long-term increase of fPSP amplitudes in both CD and HFD-fed mice (CD: n=9,5; 29.13%±9.82%; HFD: n=11,9; 37.84%±12.28%. Right: ACEA-induced fPSP increase is similar in HFD and CD-fed mice (CD: n=9,5; HFD: n=11,9; two-sample t-test p=0.59). E. Left: Schematic of whole-cell recording configuration. Middle: The CA2 PN is depolarized to induce AP firing, 25 times at 10 Hz and repeated twice, resulting in a long-term increase of PSP amplitudes in both CD and HFD-fed mice. Right: 10 Hz-induced PSP increase is similar in HFD and CD-fed mice (CD: n=10,9, 60.09%±17.40; HFD: n=9,6; 45.57%±17.06; two-sample t-test p=0.41). F. Left: Schematic of extracellular recording configuration. Middle: After prior DPDPE incubation, a 10 Hz induction protocol of 100 pulses repeated twice induces a long-term increase of SC-CA2 fPSP amplitudes in both CD and HFD-fed mice. Right: 10 Hz-induced fPSP potentiation is similar in HFD and CD-fed mice (CD: n=20,6; 35.01%±6.80%; HFD: n=15,6; 44.70%±11.91%, two-sample t-test p=0.56).

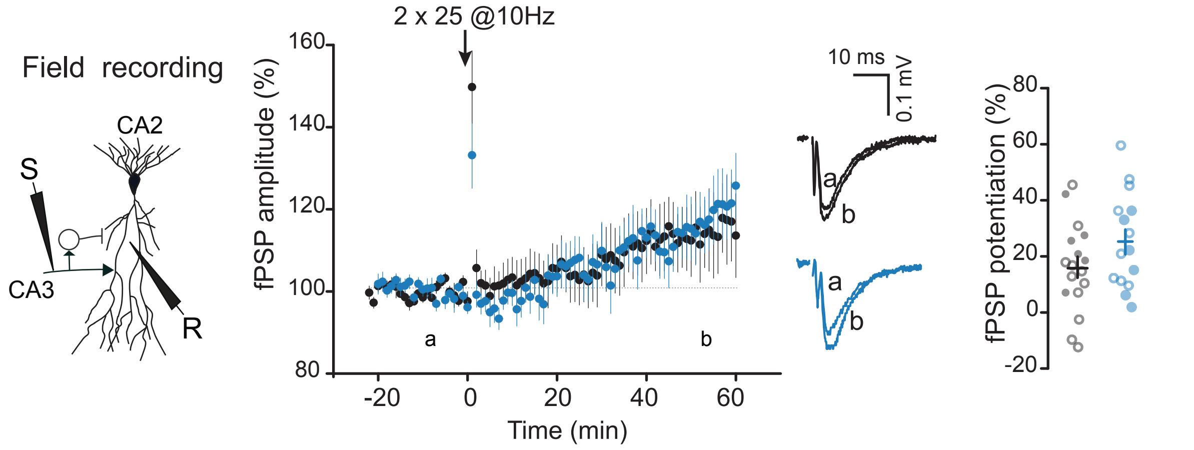


**Supplemental Figure 15.** Sub-threshold 10 Hz induction protocol. Left: Schematic of CA2 fPSP recording upon SC stimulation. Middle: Synaptic transmission was recorded before and after an induction protocol of 25 pulses of 20V stimulation at 10 Hz repeated twice, and this induced a small increase in fPSP amplitude in CA2 of both CD and HFD-fed mice without prior DPDPE incubation. Right: Long-term fPSP potentiation is significant (16.04±3.85%, one-sample t-test p= 0.0011 for CD; 25.75±4.47%, one-sample t-test p= 2.8E-4 for HFD) and similar in both CD and HFD-fed mice (CD: n=17,16; HFD n=15,14; two-sample t-test p= 0.17). Filled circles represent CA2 PNs from male mice; empty circles represent CA2 PNs from female mice.

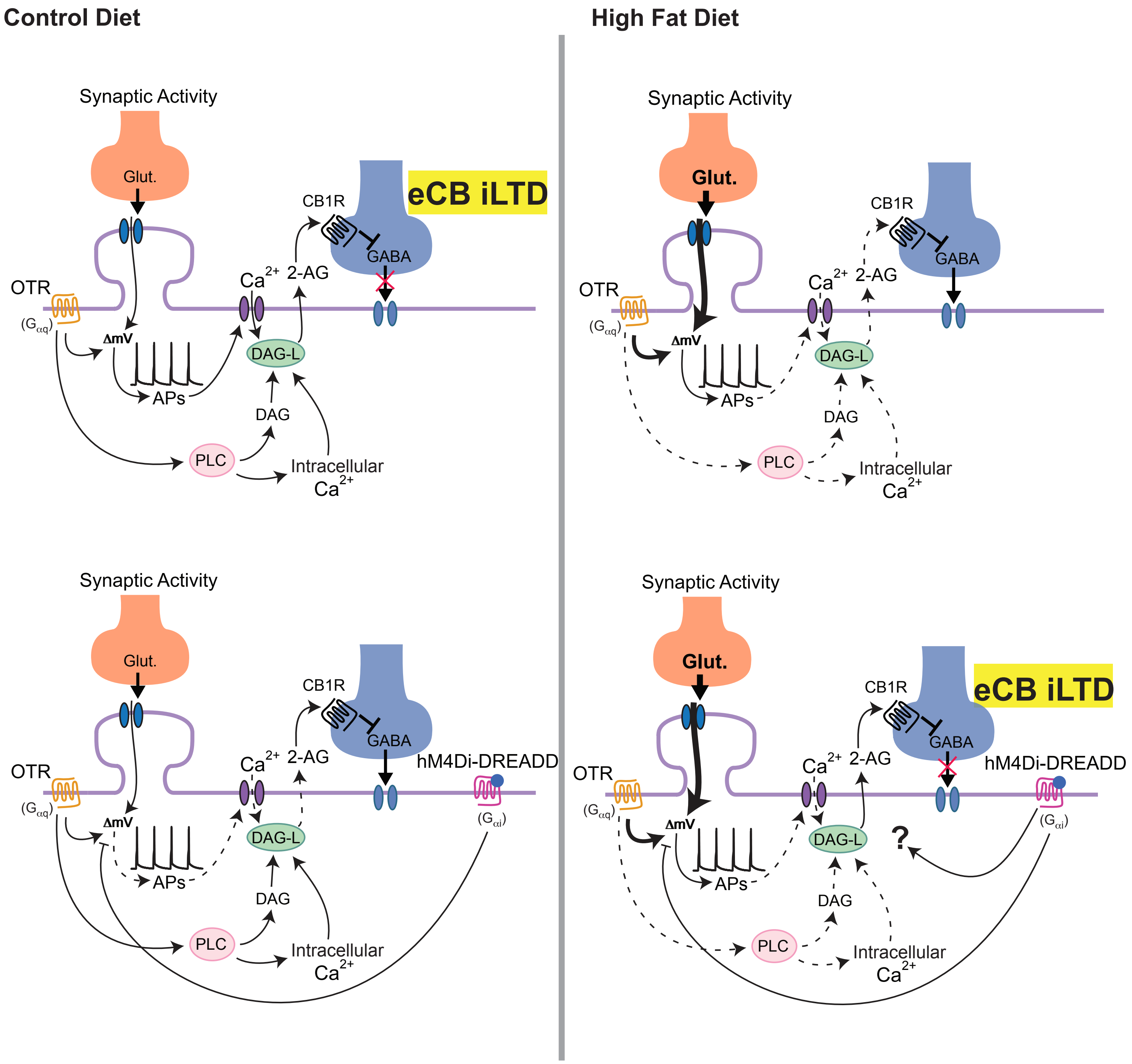


**Supplemental Figure 16.** Potential mechanisms underlying the differences in OTR signaling and eCB-mediated iLTD in CD- and HFD-fed animals and the effect of GiDREADD expression.

1) Activation of OTRs results in a larger membrane depolarization in HFD conditions (Fig 3H). However, in parallel, there are fewer OTRs detectable following social interaction in HFD-fed animals (Fig 3D) as well as less of an effect on the OTR-induced increase in synaptic transmission at 1 μM (Fig 4E). Furthermore, the decay time of action potentials is faster, resulting in less $Ca^{2+}$ influx with AP firing (Sup Fig. 11).

2) In HFD conditions, there is an increased presynaptic activity, as indicated by decreased PPR (Fig 1H). At the same time, there is a reduced increase in synaptic transmission following 1 μM OTR application (Fig 4E).

3) In HFD-treated slices, 1 μM OT application is not permissive for eCB-mediated plasticity (Fig 5E-F). This may be a result of insufficient intracellular $Ca^{2+}$. Thus, we speculate that the downstream signaling pathways following OTR activation may be ineffective in HFD conditions, as indicated by the dashed line.

4) In CD conditions, expression of the Gαi receptor hM4Di Gi-DREADD in CA2 pyramidal neurons effectively i) inhibits membrane depolarization from synaptic input, ii) reduces AP firing and iii) prevents intracellular $Ca^{2+}$ increase. Althogether, there is no eCB-mediated iLTD with synaptic activity, as established in (Loisy et al., 2022). In HFD conditions, we postulate that hM4Di receptor activation can compensate for many of the diet-induced changes in cellular physiology. With the activation of GiDREADD by CNO, 1 μM OT can promote the induction of eCB-mediated plasticity with sub-threshold stimulus in HFD conditions (Fig 6D). However, while we see rescue of eCB-mediated iLTD, the mechanistic details is a question for future studies.

Abbreviations: DAG-L, diacylglycerol lipase; PLC, Phospholipase C; DAG, diacylglycerol; OTR, oxytocin receptor; OT, oxytocin; eCB, endocannabinoid; CB1R, cannabinoid receptor type-1.
